# Functional Divergence in Gut Clostridia from Hadza Hunter-Gatherers Reveals Diet-Adapted Metabolic Machinery That Shapes Gut Ecology

**DOI:** 10.64898/2026.09.29.755216

**Authors:** Tadashi Takeuchi, Anita Reddy, Elektra K. Robinson, Kent Masakazu Kotaka, Steven K. Higginbottom, Erica D. Sonnenburg, Justin L. Sonnenburg

**Affiliations:** Department of Microbiology and Immunology, Stanford University, Stanford, CA, 94305, USA; Biophysics Program, Stanford University, Stanford, CA 94305, USA; Chan Zuckerberg Biohub, San Francisco, CA, USA; Center for Human Microbiome Studies, Stanford University School of Medicine, Stanford, CA, USA

## Abstract

Lifestyle changes associated with industrialization, including shifts in diet, are linked to extensive remodeling of the human gut microbiome. Investigating the gut microbiomes of traditional populations living non-industrialized lifestyles, such as the Hadza hunter-gatherers of Tanzania, therefore offers insight into bacterial taxa and functions depleted in industrialized microbiomes. Despite pronounced compositional differences that correspond to lifestyle, many commensal gut species are shared between industrialized and non-industrialized populations, raising a fundamental question of whether and how their functional repertoires diverge across contrasting gut environments. Here, by integrating strain-resolved metagenomics with culturomics and *in vivo* models, we identify phylogenetic and functional divergence within prevalent gut species shared between Hadza and industrialized populations, particularly among members of the Clostridia class. *Dorea longicatena*, prevalent across diverse human populations independent of lifestyle, is a poorly characterized member of Clostridia. Using strains of this *D. longicatena* isolated from either Hadza or industrialized human stool samples, we uncover metabolic pathways selectively enriched in Hadza strains, including machinery for utilizing plant-derived pectic polysaccharides. We further demonstrate that this Hadza-enriched polysaccharide-utilization machinery can be leveraged with its cognate dietary substrate to promote *D. longicatena* colonization within a model industrialized microbiota. Together, our findings reveal substantial functional divergence within gut bacterial species shared across lifestyles, providing a framework for identifying diet-responsive strains and pathways that may complement ecological and metabolic functions underrepresented in industrialized microbiotas.

## Introduction

The human gut microbiome is deeply integrated into host physiology and has been linked to health status^1,2^. Over the past decade, shotgun metagenomics has established that microbiome variation across lifestyles is accompanied by marked shifts in taxonomic composition and functional potential^3^. In particular, industrialization has been reproducibly associated with altered microbiota composition, characterized by the emergence of Bloom or Selected in Societies of Urbanization/Modernization (BloSSUM) taxa and disappearance of Volatile and/or Associated Negatively with Industrialized Societies of Humans (VANISH) taxa^4–6^, highlighting a major ecological shift in the human gut microbiome. Less attention has been paid to bacterial species shared between populations with different degrees of industrialization, even though many prevalent gut species are retained across these contrasting lifestyles^6,7^. Within these shared species, strains can show marked population-specific phylogenetic structure^7–9^, yet how this genomic variation translates into functional differences remains poorly understood. Genetic divergence at the subspecies (i.e., strain) level underlies variation in bacterial phenotypes, including polysaccharide utilization, ecological fitness within microbial communities, metabolic transformations, and effects on host immunity^10–13^. Resolving this layer of variation using a range of experimental and computational tools is essential for developing a mechanistic understanding of strain-encoded functions.

One plausible driver of such functional divergence within shared species is the human diet. Long-term dietary exposures impose sustained selective pressure on gut bacteria, particularly those encoding pathways involved in the acquisition and metabolism of complex carbohydrates^14,15^. However, relatively few studies have examined how population-associated diets shape genetic and functional diversity within bacterial species that are otherwise shared across lifestyles. One well-characterized example comes from gut *Bacteroides*, in which strains from East Asian populations encode seaweed polysaccharide degradation pathways that are rare in Western strains, consistent with cultural differences in seaweed consumption^16–18^. Building upon this finding, we previously showed in mice that the seaweed polysaccharide porphyran selectively promotes gut colonization of strains carrying the cognate polysaccharide-utilization locus (PUL)^19^. This dietary niche-enabled engraftment strategy has since been extended to humans, where dietary porphyran supported engraftment of the porphyran-utilizing species at gut densities directly proportional to the administered dose of the polysaccharide^20^. This strain was engineered to degrade dietary oxalate, highlighting that diet-linked strain functions can serve as entry points for targeted ecological manipulation and delivery of new metabolic functions in the gut.

These findings, together with the marked dietary differences between traditional and industrialized populations^21^, led us to hypothesize that industrialization is associated with alterations in metabolic repertoires among bacterial species shared across lifestyles. This model further raises the possibility that the microbiomes of traditional populations harbor functionally specialized strains that encode metabolic capacities less prevalent in industrialized populations; recovery of such functions offers a way to complement partially ablated functional repertoires of industrialized microbiotas. Testing this hypothesis, however, requires approaches that integrate strain-resolved metagenomics with experimental validation, particularly for non-model commensals whose biology remains poorly characterized.

To address lifestyle-dependent functional adaptation of gut microbes, we focused on the Hadza hunter-gatherers of northern Tanzania. Our prior work showed that Hadza gut microbiomes differ from those of industrialized populations in both taxonomic composition and predicted functional capacity. The Hadza microbiomes are characterized by many differences in composition and function, including greater representation of CAZymes associated with plant-carbohydrate utilization in Hadza than in U.S. gut metagenomes^4,22^. These patterns are consistent with the Hadza diet, which is rich in unprocessed, fiber-dense foods–a stark contrast to industrialized diets in both the composition and diversity of plant polysaccharides^21^. We therefore reasoned that the Hadza microbiome provides a unique opportunity to identify functional adaptations within bacterial species shared across populations and to determine whether the corresponding strain-encoded functions can be experimentally recovered and shown to affect gut ecology. Here, we combined strain-resolved metagenomics with experimental validation to investigate functional divergence within gut species shared between Hadza and industrialized populations.

## Results

### Hadza Clostridia are Phylogenetically and Functionally Distinct from their Non-Hadza Counterparts

To investigate functional repertoires of gut species shared between the Hadza and industrialized populations, we first sought to define the species prevalent in the Hadza and U.S., representing traditional and industrialized populations, respectively. We hypothesized that population-specific strains of a shared species would possess distinct adaptations to thrive in different gut environments. Focusing on 3,102 bacterial species identified in either the Hadza or U.S. population, we identified 123 and 113 that were present in 50% or more samples in each population, respectively. Among them, 31 of these species share 50% prevalence in both populations (Figure 1A). Notably, all 31 species belong to the Firmicutes_A phylum and the Clostridia class (Figure S1A). In the absence of a prevalence threshold, Firmicutes_A species are significantly more commonly shared between the two populations compared to the Bacteroidetes phylum, another major bacterial phylum in the human gut (23.7% of all Firmicutes_A versus 1.7% of all Bacteroidetes species, P = 7.6×10^−11^, Fisher’s test) (Figure 1B). While nearly a quarter of Firmicutes_A species are found in both of these human populations, the Bacteroidetes species exhibit nearly a complete taxonomic shift.

**Figure 1.**
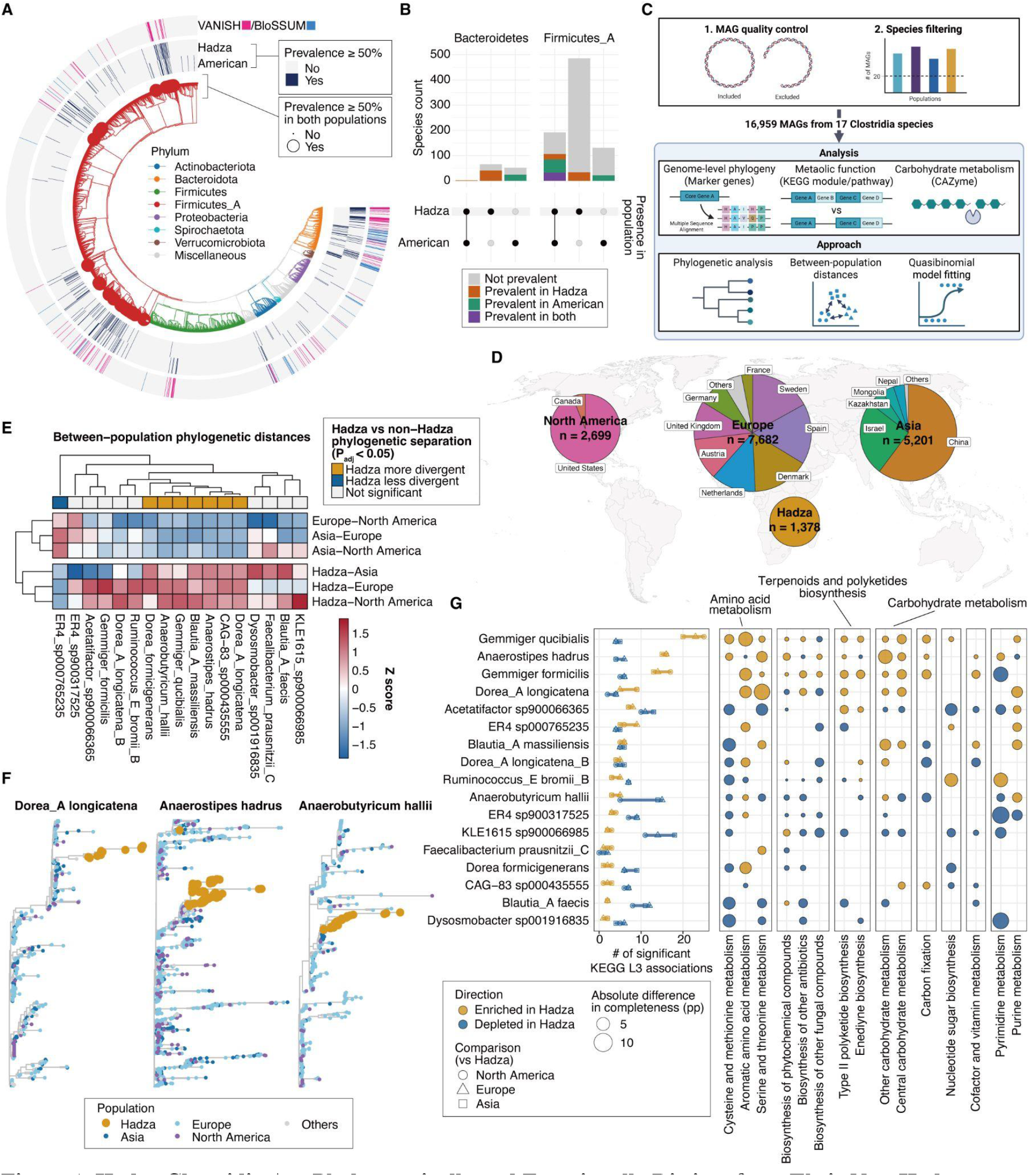
Hadza Clostridia Are Phylogenetically and Functionally Distinct from Their Non-Hadza Counterparts. (A) Phylogeny and prevalence of 3,102 SRGs identified in Hadza and U.S. metagenomes. SRGs prevalent in both populations (≥50% prevalence) are indicated by larger dots. VANISH and BloSSUM taxa are highlighted in pink and light blue, respectively, in the outer heatmap tracks. (B) UpSet plots summarizing the prevalence patterns of SRGs assigned to the Bacteroidetes and Firmicutes phyla, respectively. (C) Schematic of the genomic analysis. MAGs assigned to globally prevalent species were filtered according to genome-quality and sample-size criteria, followed by marker-gene-based phylogenetic analysis, KEGG-based functional analysis, and CAZyme analysis. (D) Pie charts showing the proportions of MAGs from four populations or geographic regions: Hadza, North America, Europe, and Asia. (E) Heatmap showing the z scores of between-population phylogenetic distances among the four populations. The top heatmap summarizes significant phylogenetic separation between the Hadza and non-Hadza populations, with green and purple indicating significantly greater and lower divergence, respectively, than distances among non-Hadza populations. (F) Marker-gene-based phylogenetic trees of representative Clostridia species exhibiting significant Hadza-biased divergence in Figure 1E, including *Dorea*_A *longicatena*, *Anaerostipes hadrus*, and *Anaerobutyricum hallii*. Dot colors indicate the population origins of the MAGs. (G) KEGG Level 3 functional differences between Hadza and non-Hadza MAGs. Numbers of Level 3 category-species associations with significantly higher or lower module completeness in Hadza MAGs (left). Points represent individual comparisons between Hadza and North America, Europe, and Asia, respectively, and horizontal lines span the minimum and maximum counts across the three comparisons. Species are ordered by their total numbers of Hadza-enriched associations. The top 15 Level 3 categories significantly associated with the largest numbers of species in the Hadza-vs-non-Hadza analysis (right). Disk size represents the absolute adjusted difference in Level 3 module completeness in percentage points (pp), and disk color indicates the direction of the association. Permutation testing (E) and quasi-binomial logistic regression (G) were used for statistical analysis. Adjusted P < 0.05 was considered significant. See also Figure S1.

We then investigated the phylogenetic and functional diversity of the 17 Clostridia prevalent in the Hadza and US (Figure 1A), including analysis of microbiome data from 24 other countries across North America, Asia, and Europe (Figures 1C and 1D). To this end, we used metagenome-assembled genomes (MAGs) available in both the Hadza study^5,6^ and the United Human Gastrointestinal Genome (UHGG) database^23^. We filtered to include species with sufficient representation in each population (≥20 MAGs per population), resulting in 16,959 MAGs taxonomically assigned to 17 Clostridia species from the four regions. All MAGs from three non-African regions were derived from industrialized and transitional populations. We further applied pre-specified QC filters and adjusted for completeness and contamination in all statistical models (See Methods).

A marker-gene-based phylogenetic analysis reveals significant population-level divergence in eight Clostridia species, with seven species showing divergence between Hadza and other populations (Figure 1E). For species showing significant phylogenetic divergence, including *Dorea*_A *longicatena*, *Anaerostipes hadrus*, and *Anaerobutyricum hallii*, Hadza MAGs clustered together to form distinct clades (Figures 1F and S1B). These findings suggest that Hadza Clostridia are phylogenetically distinct from those in other populations, a finding which is consistent with a previous global metagenome study suggesting co-diversification of humans and gut microbiomes among Clostridia species^7,8^.

To investigate functional variation among prevalent Clostridia, we next analyzed KEGG module completeness from KEGG ortholog (KO) content. The balance between Hadza-enriched and Hadza-depleted Level 3 categories varied among species. *A. hadrus*, two *Gemmiger* species, and to a lesser extent, *D. longicatena* generally showed more functional categories with significantly higher rather than lower module completeness in Hadza MAGs; whereas the majority of species showed the opposite pattern (Figure 1G; left). This overall pattern was reflected in specific functions; categories related to amino acid and carbohydrate metabolism frequently showed higher module completeness in Hadza MAGs, particularly in *Gemmiger* spp. and *D. longicatena* (Figure 1G; right). Conversely, these categories tended to show lower module completeness in Hadza MAGs from species in which Level 3 categories depleted in Hadza predominated.

Smaller but significant differences were also observed in other categories, including biosynthesis of terpenoids and polyketides, diverse small molecules that can be synthesized by bacteria. Population-associated differences in KEGG functional repertoires were also accompanied by differences in assembled MAG length within several species, suggesting that variation in genome size may contribute to the observed functional differences, though differences in assembly recovery cannot be ruled out (Figure S1C). Together, these findings reveal within-species, population-associated functional divergence among prevalent Clostridia.

To examine the ortholog-level basis of these broader functional differences, we focused on tryptophan biosynthesis and D-galacturonate degradation, which represented prominent Hadza-associated signals within amino acid and carbohydrate metabolism, respectively (Figures S1D and S1E). The majority of Hadza MAGs (i.e., different strains) from two *Gemmiger* species and *D. longicatena* encoded both subunits of tryptophan synthase, which catalyzes the final step of tryptophan biosynthesis. By contrast, these genes were significantly less prevalent in non-Hadza MAGs of the same species; K01696, in particular, was detected in less than 2% of non-Hadza MAGs (Figure S1D). This observation is consistent with a previous study reporting enrichment of tryptophan metabolic pathways in *Gemmiger qucibialis* MAGs from traditional populations^9^, suggesting that this enrichment may extend across multiple Clostridia species. Similarly, KOs constituting the complete D-galacturonate degradation module were significantly enriched in Hadza MAGs from several Clostridia species, including *D. longicatena*, *A. hadrus*, and *A. hallii* (Figure S1E). Together, these findings indicate that Hadza Clostridia are phylogenetically and functionally distinct from their counterparts in industrialized populations and suggest that industrialization is associated with non-random gains and losses in functional genes within shared bacterial lineages.

### Hadza Clostridia Possess Distinct Polysaccharide-Utilizing Repertoires from their Non-Hadza Counterparts

The enrichment of D-galacturonate degradation in several Hadza Clostridia prompted us to focus on their polysaccharide-utilization capacity. D-Galacturonate is a major component of pectic glycans, which are abundant in fruits and tubers, important components of the Hadza diet^21^. Given the important effects of dietary carbohydrates on gut bacterial composition and function, we next investigated population-associated divergence in the carbohydrate-utilization repertoires of prevalent Clostridia. CAZyme composition was substantially divergent in Hadza Clostridia, as Bray-Curtis distances between Hadza and non-Hadza population centroids were significantly greater than those among non-Hadza populations (Figure 2A). This pattern was consistent across all three Hadza–non-Hadza comparisons, with the greatest separation observed for *D. longicatena*, followed by *Acetatifactor* sp900066365, *A. hallii*, and *G. qucibialis* (Figure 2B). Significant Hadza–non-Hadza separation was detected in 16 of the 17 Clostridia species analyzed (Figure 2A), indicating the distinct CAZyme type repertoires of Hadza Clostridia. By contrast, the direction of Hadza-associated differences in CAZyme type richness was species-dependent. Total CAZyme type richness was significantly higher in Hadza MAGs from five species, including *G. qucibialis* and *D. longicatena*, but significantly lower in eight species, including *Blautia*_A *faecis* and *Acetatifactor* sp900066365 (Figure 2C). Differences in the richness of GH/PL/CE-containing CAZyme types, which are predominantly associated with glycan degradation, generally mirrored those in total CAZyme richness, whereas differences in GT-containing types, associated with glycan biosynthesis, were smaller and less consistent among species. These findings suggest that glycan-degradation capabilities are major contributors to CAZyme repertoire divergence among specific prevalent Clostridia species across human populations.

**Figure 2.**
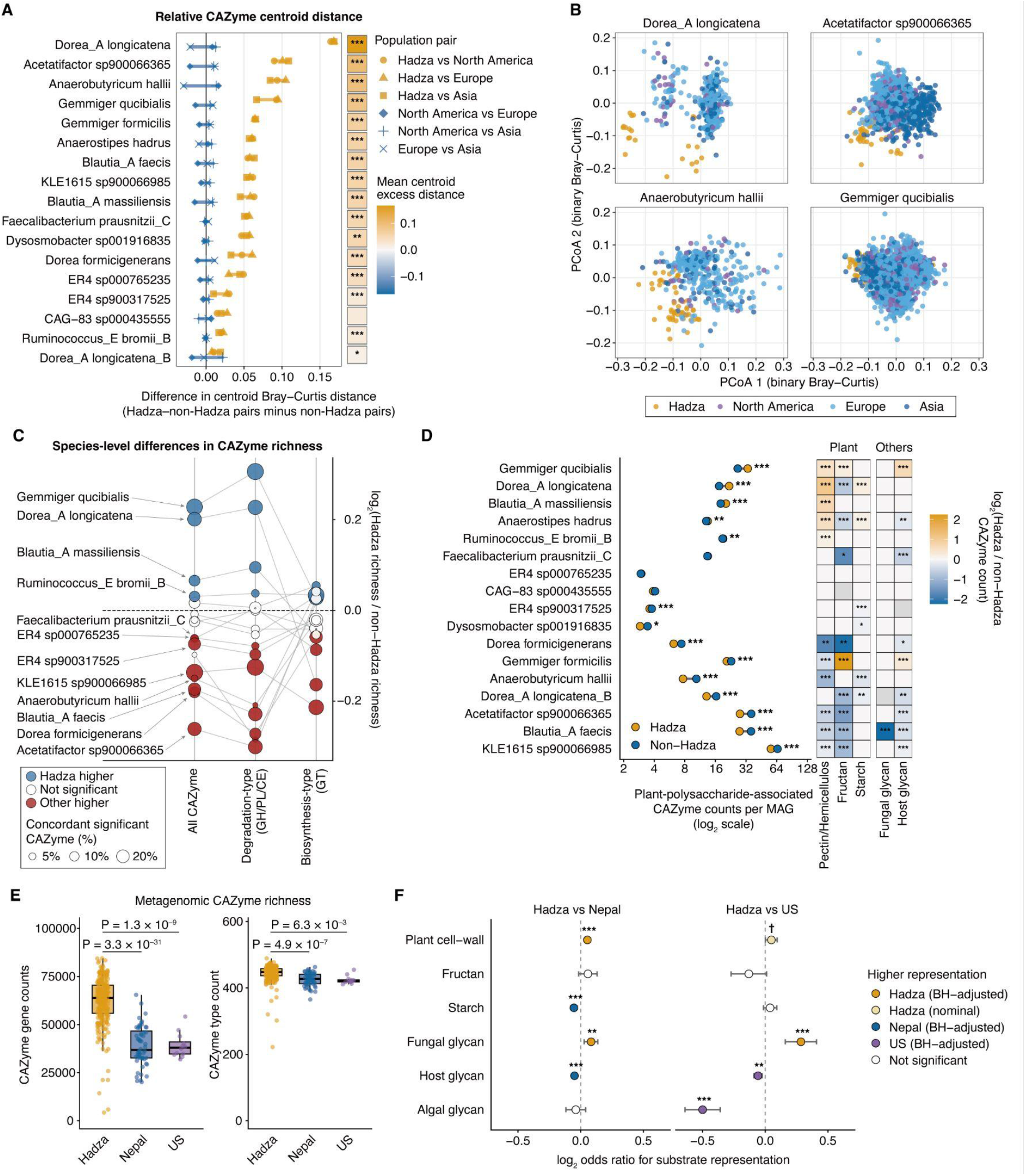
Hadza Clostridia Possess Polysaccharide-Utilization Repertoires Distinct from Those of Their Non-Hadza Counterparts. (A) Population-associated differences in CAZyme type centroids. Bray-Curtis distances were calculated between population prevalence centroids of binary CAZyme type profiles (left). Distances are shown relative to the mean of the three non-Hadza population-pair distances within each species. The heatmap shows the mean Hadza–non-Hadza distance minus the mean non-Hadza distance (right). (B) PCoA plots of Bray-Curtis distances between PCoA plots of binary Bray-Curtis distances calculated from the presence or absence of distinct CAZyme types in Hadza and non-Hadza MAGs. The results of representative Clostridia species exhibiting greater Hadza-biased divergence, including *Dorea*_A *longicatena*, *Acetatifactor* sp900066365, *Anaerobutyricum hallii*, and *Gemmiger qucibialis*, are shown. (C) Species-level differences in CAZyme richness between Hadza and non-Hadza MAGs. Points show the log_2_ ratio of adjusted richness in Hadza MAGs to the sample-size-weighted mean for MAGs from three non-Hadza populations. Lines connect estimates for all CAZymes, degradation-type (GH, PL, and CE), and biosynthesis-type (GT) within each species. Disk colors indicate significantly higher and lower richness in Hadza, and disk size indicates the fraction of tested CAZymes showing significant presence-absence differences concordant with the richness effect. (D) Adjusted counts of plant-polysaccharide-associated CAZyme genes with GH, PL, or CE annotations per MAG in Hadza and non-Hadza groups (left). Plant-polysaccharide counts comprise genes associated with plant cell-wall-, fructan-, and starch-associated glycans. Orange and blue points indicate Hadza and weighted non-Hadza estimates, respectively. The heatmap (right) shows log_2_ ratios of adjusted gene counts for individual substrate classes. Orange and blue indicate significantly higher and lower counts in Hadza, respectively; white indicates tested but nonsignificant comparisons, and gray indicates comparisons excluded by QC. Species are ordered by the adjusted count difference between Hadza and weighted non-Hadza estimates. (E) Richness of detected CAZyme genes and CAZyme types in rarefied fecal metagenomes from Hadza, Nepali, and U.S. participants (n = 249, 56, and 12). (F) Population-associated enrichment of putative polysaccharide-degrading CAZyme types associated with broad substrate classes. Points and error bars show log_2_ odds ratios and 95% confidence intervals comparing Hadza with Nepali or U.S. metagenomes. Dot colors indicate the population with the higher proportion, with dark and light fills indicating associations meeting FDR-adjusted and nominal significance thresholds, respectively. Permutation testing (A), quasi-Poisson regression with a log link (C–E), and bias-reduced binomial logistic regression (C and F) were used for statistical analysis. *adjusted P < 0.05; **adjusted P < 0.01; ***adjusted P < 0.001; † P < 0.05. See also Figure S2.

To investigate the functional aspects of CAZymes in Hadza Clostridia, we next analyzed their predicted substrate specificities. Although we initially expected Hadza Clostridia to harbor universally higher numbers of CAZymes related to plant polysaccharides, given the high consumption of plant fiber by the Hadza^21^, the direction of these differences again varied among species (Figure 2D). Focusing on those CAZyme subsets, we found that *G. qucibialis* and *D. longicatena* had higher numbers of CAZyme gene copies per MAG in the Hadza population, whereas *B. faecis* and *Acetatifactor* sp900066365 had lower numbers, consistent with the overall CAZyme richness analysis. At the substrate level, differences in total plant-polysaccharide CAZyme count generally paralleled the pattern for plant cell-wall glycans, including pectin, hemicellulose, and cellulose. By contrast, CAZymes associated with fructan tended to be depleted in Hadza MAGs, even in species showing Hadza enrichment of total CAZymes related to plant polysaccharides, including *D. longicatena* and *A. hadrus*. Consistently, GH32, a CAZyme family involved in fructan degradation, was significantly underrepresented in Hadza MAGs from eight Clostridia species but overrepresented in only two, both belonging to *Gemmiger* (Figure S2A). These findings suggest that population-associated differences in carbohydrate-utilization repertoires vary across both substrate classes and species.

To determine whether population-associated differences in CAZyme repertoires extended beyond prevalent Clostridia species, we additionally analyzed CAZymes in Hadza fecal metagenomes, using U.S. and Nepali metagenomes as representatives of industrialized and transitional populations, respectively^6^. Consistent with a previous study^4^, both CAZyme gene richness and CAZyme type richness were significantly higher in rarefied Hadza metagenomes (Figure 2E). The directions of population-associated differences in CAZyme type detection were concordant between the two comparisons, with Hadza–U.S. and Hadza–Nepali effect sizes positively correlated (Pearson’s R = 0.67, P < 2.2 × 10^−16^; Figure S2B). This finding suggests consistent differentiation of the Hadza CAZyme repertoire relative to both industrialized and non-industrialized non-Hadza populations. Substrate-level analysis further revealed that putative polysaccharide-degrading CAZyme types associated with plant cell-wall glycans were enriched in Hadza metagenomes, with significant enrichment in the Hadza–Nepali comparison and nominal increase (i.e., unadjusted P < 0.05) in the Hadza–U.S. comparison (Figure 2F). By contrast, CAZyme types associated with other plant-derived glycans, such as fructan and starch, showed either no difference or depletion in Hadza metagenomes. CAZyme types associated with fungal polysaccharides were enriched in Hadza metagenomes, whereas those associated with host glycans were depleted in both comparisons. Together, these findings extend the strain-resolved observations to the community level, suggesting distinct polysaccharide substrate preferences in the Hadza gut microbiome, particularly among plant-derived polysaccharides, consistent with inter-population differences in quantity and types of plants consumed.

### Hadza *D. longicatena* Isolates Possess a Polysaccharide-Utilizing Machinery for Arabinan

Our MAG analysis reveals that particular Clostridia show significant phylogenetic and functional divergence between Hadza and non-traditional populations. *D. longicatena* and *G. qucibialis* are among the most promising candidate species showing consistent phylogenetic divergence and functional enrichment in Hadza MAGs, including CAZyme diversity. We therefore sought to characterize aspects of these species’ functionality by isolating strains from Hadza fecal samples. Using 21 fecal samples and a culturomics approach, we isolated 737 strains (see Methods). Among them, we identified eight strains of *D. longicatena*, taxonomically assigned by Sanger sequencing of the 16S rRNA gene. By whole-genome sequencing, we further identified that five were classified as *Dorea*_A *longicatena* and two strains as *Dorea*_A *longicatena*_B, with three *Dorea*_A *longicatena* strains isolated from the same subject being nearly identical (Figure 3A). On the other hand, one isolate was classified as *Dorea*_A sp934294775, which is phylogenetically close to *Dorea*_A *longicatena* but was only recently identified in the Hadza metagenome study^6^. We confirmed that the prevalence of *Dorea*_A sp934294775 was 55.7% in the Hadza but zero in the U.S. population, suggesting this is a VANISH species (Figure S3A). Unlike *D. longicatena*, we could not isolate *G. qucibialis* strains, prompting our focus on *D. longicatena* to investigate Hadza-specific functions in Clostridia.

**Figure 3.**
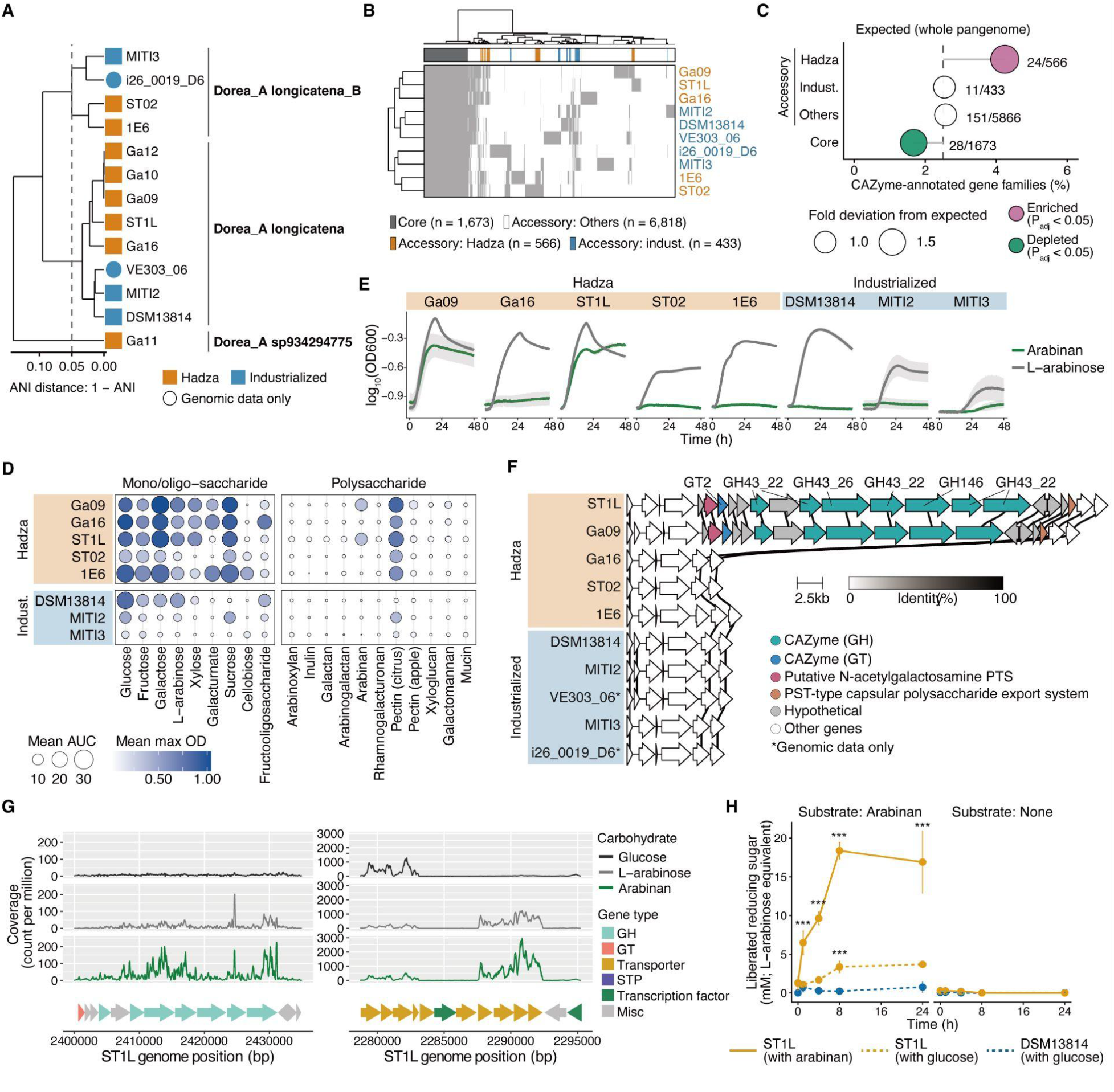
Hadza *D. longicatena* Isolates Possess Arabinan-Utilization Machinery. (A) ANI-based dendrogram of *Dorea* isolates from Hadza and industrialized populations, including the type strain DSM 13814. Two complete genome assemblies from NCBI were also included as industrialized-population genomes. (B) Pangenome heatmap of *D. longicatena* isolates. The top bar indicates the prevalence of each gene family among the isolates. Gene families were defined as core when detected in all isolates and as population-enriched accessory when detected in at least two isolates from one population and absent from all isolates from the other population. (C) Proportions of CAZyme-annotated gene families among four gene-prevalence categories. The proportion in each category was compared with the genome-wide expected proportion. Significant enrichment relative to the expected proportion is indicated in pink. (D) Bubble plot summarizing the growth metrics of eight *D. longicatena* isolates cultured in depleted medium supplemented with 0.5% of the indicated carbohydrate as the sole carbon source. Disk size and color indicate the AUC and maximum OD, respectively. (E) Growth curves of eight *D. longicatena* isolates cultured in depleted medium supplemented with 0.5% arabinan. Shaded areas indicate the standard deviation (n = 3 per group). (F) Alignment of the GH43/GH146 gene cluster in Hadza *D. longicatena* strain ST1L (ST1L_wgs_02240–ST1L_wgs_02261) with homologous regions in nine *D. longicatena* isolate genomes from Hadza and industrialized populations, including two complete genome assemblies from NCBI. (G) RNA-sequencing analysis showing the transcriptional activity of the GH43/GH146 cluster (left) and the putative L-arabinose transporter cluster (right) in Hadza *D.* (H) Reducing sugars released by resting cells of Hadza *D. longicatena* strain ST1L and the type strain DSM 13814 during incubation in PBS containing 0.5% arabinan or in Fisher’s exact test (C) and mixed-design repeated-measures ANOVA followed by pairwise comparisons of estimated marginal means with Holm adjustment (H) were used for statistical analysis. Asterisks denote comparisons between baseline and each subsequent time point (H). ***adjusted P < 0.001. See also Figure S3.

Our analysis included five phylogenetically independent Hadza *D. longicatena* strains plus three isolates from industrialized populations. In addition, we utilized complete assemblies of two additional industrialized sample-derived isolates available on NCBI. Using these 10 isolate genomes, we first conducted a comparative genomic analysis to understand the characteristics of Hadza-enriched gene families. Focusing on genes detected in at least two genomes from one population but absent from all genomes in the other, we identified 566 and 433 genes specific to the Hadza and industrialized populations, respectively (Figure 3B). With Clusters of Orthologous Groups (COG) annotations, we found that most major metabolic COG classes were disproportionately represented in the core genome, consistent with their roles in conserved cellular metabolism (Figure S3B). By contrast, carbohydrate transport and metabolism genes (COG category G) were overrepresented in the Hadza-restricted accessory cluster, indicating that carbohydrate utilization represents a major axis of population-associated strain diversification. Similarly, CAZyme annotations were significantly enriched among Hadza-restricted clusters relative to their expected pangenome-wide frequency (Figure 3C). Together with our MAG-based analysis (Figure 2E), these findings reveal that the Hadza-associated functional repertoire in *D. longicatena* is particularly enriched in polysaccharide-utilizing machinery.

Next, we performed growth assays to ask whether Hadza *D. longicatena* isolates show divergent carbohydrate utilization. Overall, Hadza isolates showed higher area under the curves (AUCs) of growth than industrialized counterparts with mono- and oligosaccharides, reflecting their basic carbohydrate metabolism (Figure 3D and S3C). Specifically, galacturonate, a primary monomer of pectin chain, was exclusively utilized by three Hadza isolates (Figure S3C), consistent with the enrichment of the galacturonic acid-degrading pathway among Hadza Clostridia MAGs (Figure S1E). By contrast, with fructooligosaccharides, a shorter fructan, Hadza isolates, except Ga16, achieved approximately half of the maximum growth expected with its monomer fructose (Figure S3D). Their CAZyme profile further revealed that Hadza isolates possess significantly fewer copies of GH32, a fructan-degrading CAZymes (Figures S3E and S3F), consistent with depleted GH32 among many Hadza Clostridia, including *D. longicatena* (Figure S2A). Furthermore, none of the Hadza isolates possessed GH172, a GH family implicated in the degradation of caramelized fructose-containing oligosaccharides derived from heated sugars in ultra-processed foods (Figure S3F)^24^.

Unlike mono- and oligosaccharides, only a few polysaccharides were utilized by *D. longicatena*, suggesting variability in depolymerization and utilization capacity. Whereas pectin from citrus was utilized by most isolates regardless of origin, pectin from apple was utilized poorly, consistent with specificity of the strains for a subset of the structurally diverse pectins. Arabinan, a neutral sugar side chain attached to pectin, was exclusively utilized by two Hadza *D. longicatena* isolates (Figures 3D and 3E). This is despite all isolates showing the abilities to utilize L-arabinose, the monomer, suggesting that CAZymes that degrade arabinan, rather than L-arabinose transporters and downstream catabolic pathways, are specific to these two Hadza isolates. None of the *D. longicatena* isolates utilized other pectic polysaccharides, such as galactan or arabinogalactan.

We further identified a gene cluster that contains members of GH43 subfamilies (GH43_22 and GH43_26) and GH146, which encode L-arabinofuranosidases known to degrade arabinan^25^, exclusively in the two Hadza *D. longicatena* isolates that utilize arabinan (Figures 3F and S3E). This gene cluster was significantly prevalent in Hadza *Dorea*_A *longicatena* MAGs compared to >230 non-Hadza MAGs analyzed (Figure S3G). Notably, Fiji-derived genomes showed intermediate prevalence of similar gene clusters, implying an association with non-industrialized lifestyles. RNA sequencing confirmed transcription of the putative GH43/GH146 cluster in arabinan-containing medium, whereas transcription remained low under glucose or L-arabinose conditions (Figure 3G). However, unlike typical *Bacteroides* PULs, in which transporters are encoded in proximity to cognate CAZymes^25^, this putative arabinan gene cluster does not contain transporters. Instead, we identified another gene cluster containing multiple putative arabinose transporters, located approximately 120 kbp upstream of the GH43/GH146 cluster (Figure 3G). These putative arabinose transporters were active in both arabinan and L-arabinose-containing media, but not in the glucose condition, supporting their involvement in arabinan utilization.

The separation of CAZyme and transporter loci suggests a model in which GH43 enzymes function at the bacterial cell surface, as described for Gram-positive bacteria that lack a periplasmic compartment^26^. In this model, surface-associated CAZymes cleave arabinan outside the cytoplasm, generating mono- and/or oligosaccharides that are subsequently imported by transporters. Supporting this model, growth-arrested ST1L cells pre-cultured in arabinan-containing medium produced a strong arabinan-dependent increase in reducing-end signal over 24 h, consistent with robust arabinan hydrolysis activity (Figure 3H). Notably, ST1L cells pre-cultured in glucose-containing medium showed only 21.8% of the reducing-end signal observed in arabinan-pre-cultured ST1L cells, suggesting that this hydrolysis activity is inducible. In contrast, DSM 13814, which lacks the GH43/GH146 cluster, did not show an arabinan-dependent increase in reducing-end signal. Together, these findings reveal a distinct arabinan-utilization machinery that is predominantly conserved in Hadza *D. longicatena*.

### Hadza *D. longicatena* Shares a Putative PUL with a Phylogenetically Close Hadza-Specific Species

In addition to the GH43/GH146 cluster, we found other CAZymes specific to Hadza *D. longicatena* isolates, such as GH43_11, GH5_44, and GH39 (Figure S3E). These Hadza-specific CAZymes were encoded in close proximity, forming another putative PUL in Hadza *D. longicatena* ST1L (Figure 4A). Notably, we further identified a similarly aligned PUL in the Hadza-specific *Dorea* strain Ga11, taxonomically assigned to *Dorea*_A sp934294775 (Figures 4A and S3A). Other Hadza *D. longicatena* isolates, such as ST02, Ga16, and 1E6, possessed only the proximal several genes of the PUL, whereas no industrialized isolates possessed it. Although GH43_11, GH5_44, and GH39 were expected to degrade pectic polysaccharides, commercially available pectin did not induce transcription of genes in this PUL (Figure S4A), leaving its target substrate unknown. Given the structural diversity of polysaccharides in edible plants and the limited range of structures represented by commercially available fiber preparations^27^, one possibility is that this PUL targets a polysaccharide present in the Hadza diet that is not represented by commercially available substrates. In line with this hypothesis, the polysaccharide-utilizing genes in this PUL, especially GH43_11, GH5_44, and their adjacent genes (Ga11_wgs_01250 to 01253), were shared broadly and almost exclusively among Hadza *Dorea* spp., including two subspecies of *D. longicatena* and *Dorea*_A sp934294775 (Figure 4B), suggesting a biological role associated with the Hadza gut environment.

**Figure 4.**
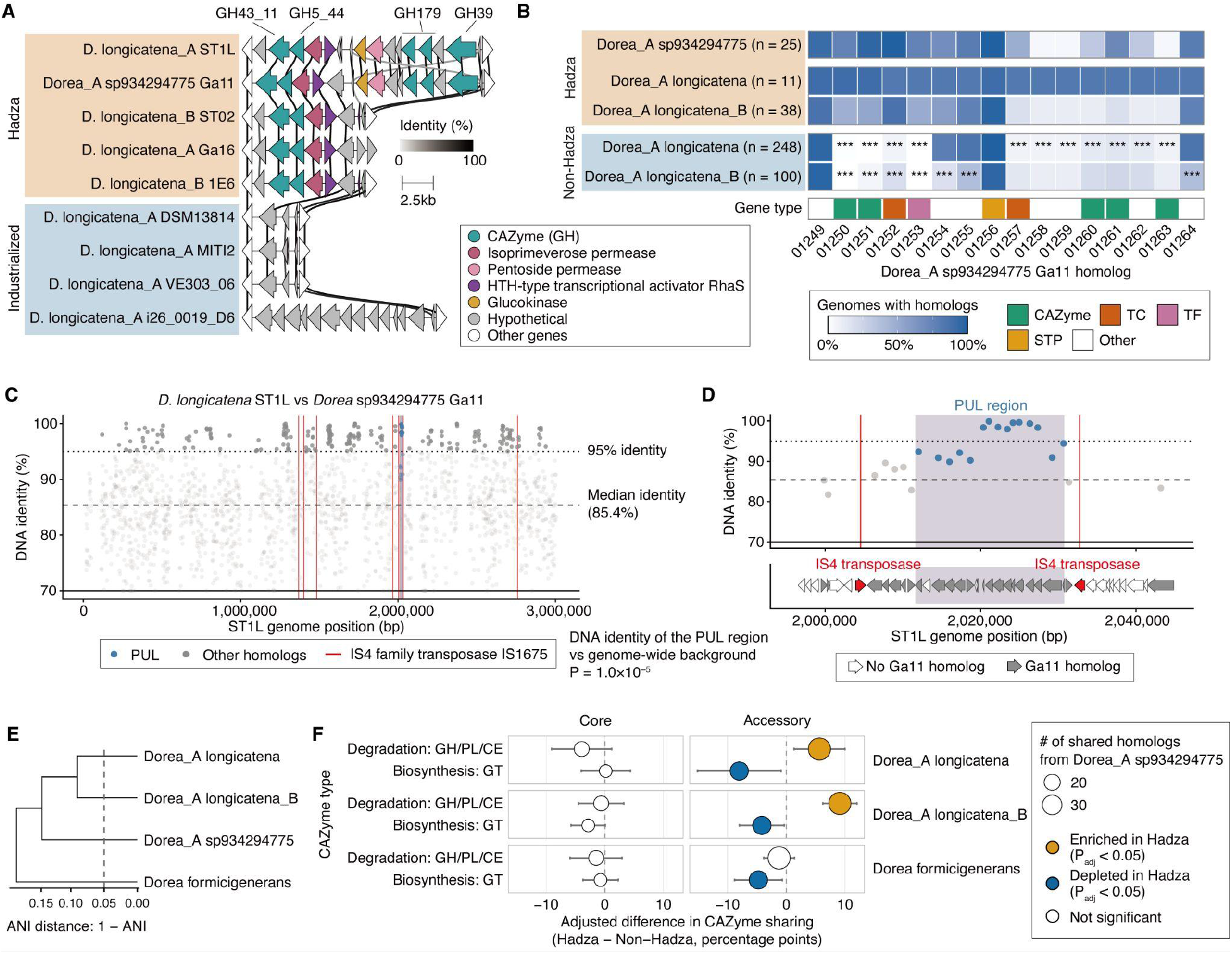
Hadza *D. longicatena* Shares a Putative PUL with a Phylogenetically Related Hadza-Specific Species. (A) Alignment of the GH43/GH5 PUL in Hadza *D. longicatena* strain ST1L (ST1L_wgs_01887–ST1L_wgs_01905) with homologous regions in Hadza *Dorea*_A sp934294775 strain Ga11 and six *D. longicatena* isolate genomes from Hadza and industrialized populations. (B) Local sequence conservation of homologs from Hadza *Dorea*_A sp934294775 strain Ga11 in *D. longicatena* and *D. longicatena*_B genomes from Hadza and industrialized populations. The white-to-blue color scale indicates homolog prevalence. Asterisks denote significant differences between the Hadza and industrialized populations within each taxonomic group. (C and D) Genome-wide map of Hadza *D. longicatena* strain ST1L showing the nucleotide identities of homologs shared with Hadza *Dorea*_A sp934294775 strain Ga11 (C) and a magnified view of the GH43/GH5 PUL (D). Homologs with ≥95% nucleotide identity are highlighted. Red lines indicate the locations of the IS4-family transposase IS1675, and blue dots indicate homologs from the GH43/GH5 PUL. Gray and white gene arrows in (D) indicate homologs shared with Ga11. (E) ANI-based dendrogram of SRGs from four *Dorea* species. (F) Sharing of CAZyme-annotated homologs between Hadza *Dorea*_A sp934294775 genomes and Hadza or industrialized genomes from three *Dorea* species. CAZymes are classified as degradation-related or biosynthesis-related. Adjusted differences in prevalence between Hadza and industrialized genomes are shown. Fisher’s exact test (B), permutation test, and linear regression (F) were used for statistical analysis. ***adjusted P < 0.001. See also Figure S4.

We next investigated the evolutionary relationship of the Hadza-associated GH43/GH5 PUL by comparing the corresponding genomic regions of *D. longicatena* ST1L and *Dorea*_A sp934294775 Ga11. Homologous genes within the PUL showed markedly elevated nucleotide identity, with many homologs sharing >95% identity and several approaching 99% identity; the median nucleotide identity within the PUL was significantly higher than the genome-wide background (P = 1.0 × 10^−5^; Figures 4C and 4D). The PUL also formed a localized region of dense gene-level homology between ST1L and Ga11, whereas the 10-kb genomic regions immediately surrounding the PUL contained multiple genes without detectable Ga11 homologs (Figure 4D; lower panel).

This unusually high sequence conservation between two different *Dorea* species suggests that this PUL may have been horizontally transferred between these lineages or similarly acquired from a closely related common donor. Supporting this possibility, we identified seven highly similar IS4-family transposases in the ST1L genome (Figures 4C and S4B), with two 100%-identical, oppositely oriented copies positioned on opposite sides of the PUL-containing region (Figure 4D). We further identified two 1,691-bp IS-like copies sharing 99.4% nucleotide identity, with corresponding imperfect terminal inverted repeats (TIRs) (Figures S4C and S4D), suggesting a composite-transposon-like architecture surrounding the PUL-containing region. Together with the localized elevation in sequence identity and gene homology between ST1L and Ga11 (Figures 4C and 4D), these structural features provide additional evidence consistent with horizontal transfer of the PUL-containing. By contrast, strains containing only the proximal portion of the PUL, such as Ga16, ST02, and 1E6, showed mixed mobile-element contexts. Whereas Ga16 contained several heterogeneous transposases nearby, no obvious PUL-associated transposases were detected in ST02 or 1E6 (Figure S4E). These findings suggest lineage-specific retention and genomic remodeling of the locus, including partial retention or truncation of the PUL in some strains.

To investigate whether gene sharing between the Hadza-associated VANISH *Dorea* and *D. longicatena* extends beyond the GH43/GH5 PUL, we analyzed the representation of *Dorea*_A sp934294775 homologs in MAGs from three *Dorea* species that are prevalent in both Hadza and non-Hadza populations (Figures 4E and S1A). Genome-wide sharing of *Dorea*_A sp934294775 accessory homologs was modestly but significantly elevated in Hadza *Dorea*_A *longicatena* compared with non-Hadza strains (adjusted P = 0.018), whereas a similar but non-significant trend was observed in *Dorea*_A *longicatena*_B (adjusted P = 0.10) (Figure S4F). There was no difference in *Dorea formicigenerans*, a more distant *Dorea* species. By contrast, *Dorea*_A sp934294775 homologs that are highly prevalent across MAGs (≥90%) showed no population-associated difference in either *Dorea*_A *longicatena* lineage, while a modest increase was observed in non-Hadza *D. formicigenerans* (Figure S4F). We next asked whether population-associated accessory sharing was concentrated in carbohydrate-active functions. Among accessory CAZyme homologs detected in *Dorea*_A sp934294775, those involved in glycan degradation, including GH, PL, CE, and CBM, showed significantly greater adjusted sharing with Hadza than non-Hadza genomes in both *Dorea*_A *longicatena* species, whereas no corresponding Hadza enrichment was observed in *D. formicigenerans* (Figure 4F). By contrast, accessory GT homologs, involved in glycan biosynthesis, were preferentially represented in non-Hadza genomes across three *Dorea* lineages. No significant population-associated differences were observed for either CAZyme class among high-prevalence CAZyme homologs. Together, these findings suggest preferential sharing of glycan-related accessory genes between the Hadza-associated VANISH lineage and closely related Hadza *D. longicatena* lineages, potentially contributing to population-associated diversification of polysaccharide-utilization repertoires.

### Polysaccharide Utilization Machinery in Hadza *D. longicatena* Supports Competitive Gut Colonization

Unlike *Bacteroides* spp., many Lachnospiraceae species, including *D. longicatena*, colonize poorly in experimental gut communities in mice and show limited engraftment following fecal microbiota transplantation in humans^28,29^. Therefore, robust and reproducible strategies for gut engraftment of these bacteria are needed to investigate and realize the potential of their metabolic functions, including metabolite production. We hypothesized that the arabinan utilization machinery enriched in Hadza-associated *D. longicatena* provides access to a dietary niche in the presence of arabinan, supporting stable colonization within a complex bacterial community. Previous work in our lab has demonstrated the ability of the seaweed polysaccharide porphyran to create a privileged niche enabling engraftment of a cognate utilizing strain^19,20^. Here we investigate the utility of niche creation, without complete exclusivity to the incoming strain that porphyran afforded. To test our hypothesis, we used a synthetic community comprising 14 representative human gut species from industrialized populations (14SM) as the background microbiota^30,31^. All six *Bacteroides* species showed moderate to robust growth in monoculture with arabinan as the sole carbon source (Figure S5A).

Using this 14SM community and two *D. longicatena* strains, Hadza ST1L and the type strain DSM 13814, representing an arabinan utilizer and non-utilizer, respectively, we next tested whether arabinan specifically increases the abundance of ST1L *in vitro*. Principal coordinate analysis (PCoA) shows significant bacterial community alterations largely driven by carbohydrate conditions (Figure 5A). In addition, the significant carbohydrate-by-*D. longicatena* strain interaction indicates that the effect of *D. longicatena* supplementation on community composition differs among carbohydrate conditions (PERMANOVA, R^2^ = 0.039, P = 3.5×10^−3^). We further observed significant differences in inter-community Bray-Curtis distances among three carbohydrate conditions, in which arabinan showed the highest separation between two communities (Figure S5B). These findings suggest a synergistic effect of ST1L and arabinan on the bacterial community composition. As expected, ST1L showed significantly higher relative abundance in cultures with arabinan compared to DSM 13814, occupying the highest abundance among the community members (Figures 5B and 5C). Consequently, three species showed a significant reduction in relative abundance in the 14SM + ST1L community. Nevertheless, ST1L also increased significantly under the glucose condition, although to a lesser extent than under the arabinan condition. The relative abundance of ST1L and DSM13814 was comparable in cultures with L-arabinose, albeit their abundance was high (up to 50%). Notably, arabinan increased a subset of *Bacteroides* species, such as *B. cellulosilyticus*, *B. vulgatus*, and *B. uniformis*, while limiting the abundance of other *Bacteroides*, even though all the *Bacteroides* species were shown to utilize arabinan in single cultures (Figures S5A and S5C). These findings suggest a robust functional relationship between arabinan and Hadza *D. longicatena* ST1L even in the context of several arabinan-utilizing community members.

**Figure 5.**
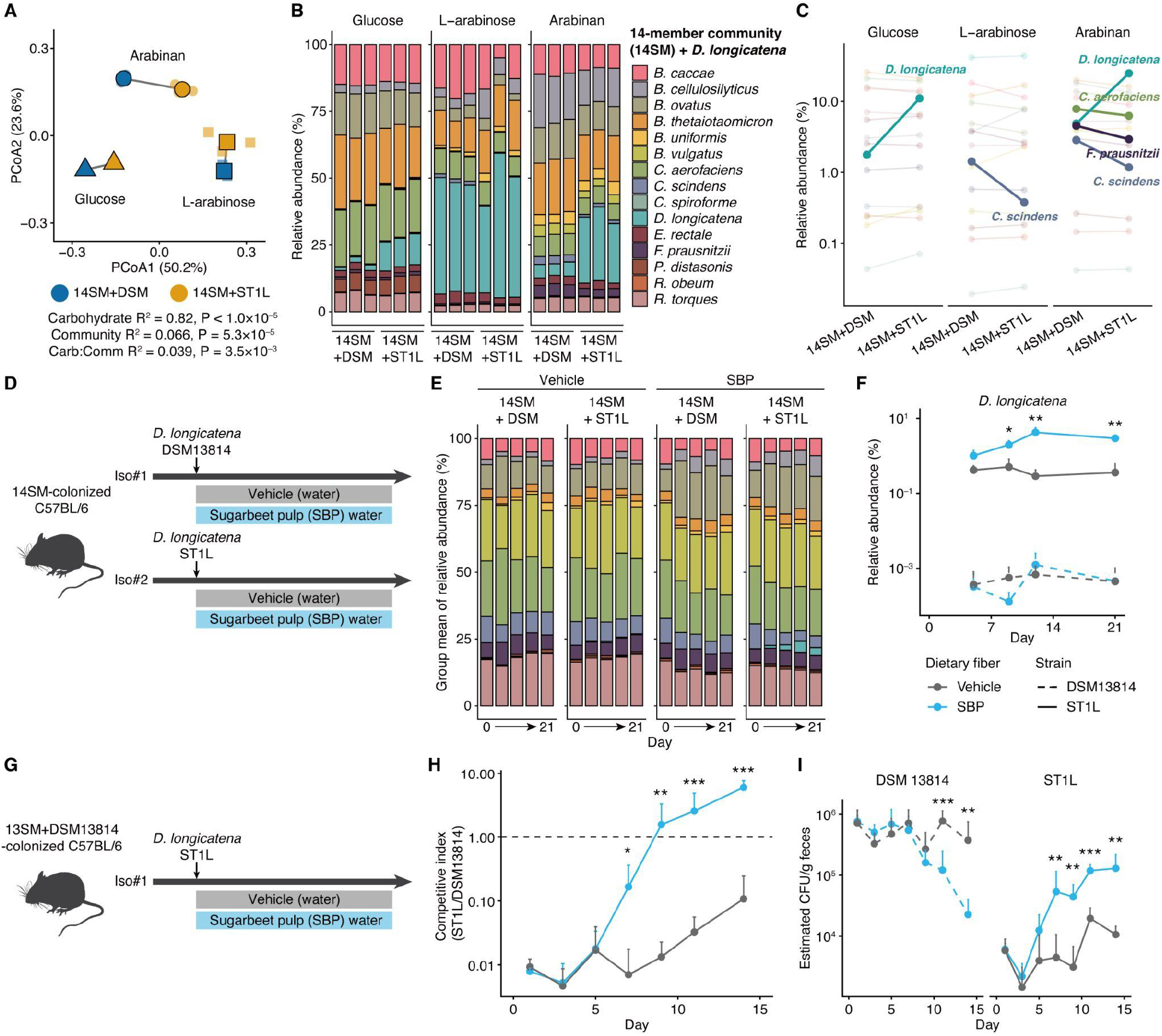
Hadza-Enriched Polysaccharide-Utilization Machinery in *D. longicatena* Supports Competitive Gut Colonization. (A) PCoA plot of Bray-Curtis distances among *in vitro* communities comprising the 14-member synthetic community (14SM) plus Hadza *D. longicatena* strain ST1L or the type strain DSM 13814. Depleted medium supplemented with 1% glucose, L-arabinose, or arabinan was used. Larger symbols connected by solid lines indicate the centroids of the community-carbohydrate groups. PERMANOVA results are shown at the bottom (n = 3 per group). (B and C) Relative abundances of the 14SM members and *D. longicatena* strains in the *in vitro* communities (B). Community members exhibiting significant differences in relative abundance between the two *D. longicatena*-harboring communities are highlighted in (C). (D) Design of a gnotobiotic experiment testing the effect of SBP on the colonization of Hadza *D. longicatena* strain ST1L in mice harboring the 14SM. (E) Relative abundances of the 14SM members and *D. longicatena* strains in the gnotobiotic mice. (F) Relative abundance of *D. longicatena* strains in the gnotobiotic mice. Asterisks denote significant differences between ST1L-colonized mice administered SBP and those administered vehicle (water) at each time point. (G) Design of a gnotobiotic experiment testing the effect of SBP on the colonization of Hadza *D. longicatena* strain ST1L in mice harboring the 13SM plus the *D. longicatena* type strain DSM 13814. *C. scindens* was removed from the 14SM to allow robust gut colonization by DSM 13814. (H) Competitive index of ST1L, expressed as the ratio of the relative abundance of ST1L to that of DSM 13814. (I) Estimated absolute abundances of ST1L and DSM 13814 in the gnotobiotic mice, expressed as CFU equivalents. Permutational ANOVA (A), two-sided Welch’s t-test (C), and mixed-design repeated-measures ANOVA of log_10_-transformed relative abundances followed by pairwise comparisons of estimated marginal means with Holm adjustment (F, H, and J) were used for statistical analysis. Asterisks denote comparisons between the vehicle and SBP groups within each strain group (F, H, and J). *adjusted P < 0.05; **adjusted P < 0.01; ***adjusted P < 0.001. See also Figure S5.

Next, using the same 14SM community, we investigated *D. longicatena* colonization in a gnotobiotic mouse model (Figure 5D). Sugar beet pulp (SBP), a dietary fiber source enriched in arabinan and confirmed to support ST1L growth *in vitro*, was used for the animal studies (Figure S5D). Consistent with our *in vitro* findings, SBP was the dominant factor explaining variation in microbiota composition, particularly at later time points (Figure S5E). Accordingly, dietary SBP was associated with changes in the relative abundance of multiple species in both ST1L and DSM 13814 communities (Figures S5F and S5G). Nevertheless, SBP increased the relative abundance of ST1L by up to 10-fold, an effect not observed in mice colonized with DSM 13814, which remained at significantly lower relative abundance regardless of SBP administration (Figures 5E and 5F). Although several taxa showed significant but small SBP effects that were not reproduced across both communities, a significant overall SBP by strain interaction was detected only for *D. longicatena* (Figures S5F and S5G). Together, these findings demonstrate that SBP selectively enhances ST1L colonization within a complex gut community, consistent with the arabinan-dependent growth phenotype observed *in vitro*.

We next asked whether ST1L exhibits a competitive advantage in mice previously colonized with DSM 13814 when SBP is administered (Figure 5G). Because DSM 13814 colonization is inhibited by *C. scindens*, a member of the 14SM community, and *C. scindens* abundance was strongly negatively correlated with ST1L abundance in the present study (Figure S5H), we removed *C. scindens* to generate a 13SM community that supported robust baseline colonization by DSM 13814 (Figure S5I). Despite the prior colonization of DSM 13814, ST1L rapidly increased in abundance following SBP administration and surpassed DSM 13814 within two weeks (Figure 5H). ST1L also gradually increased without SBP but remained approximately 1 log_10 u_nit less abundant than DSM 13814, consistent with the inhibitory priority effect of a first arriving cognate strain reported in other species^19,32,33^. More strikingly, the absolute abundance of DSM 13814 declined rapidly following SBP administration but remained stable in its absence, suggesting that the SBP-supported expansion of ST1L enabled it to compete with and displace the previously colonized DSM 13814 population (Figure 5I). Together, these findings indicate that the Hadza-enriched polysaccharide utilization machinery confers a competitive advantage on ST1L within a gut community, even in the presence of an industrialized *D. longicatena* strain.

## Discussion

Using strain-resolved metagenomics and culturomics, we identify previously unrecognized functional divergence between Hadza and industrialized gut bacteria encoded at the strain level, particularly among globally prevalent Clostridia species. Focusing on *D. longicatena*, we demonstrate that Hadza-enriched metabolic machinery enables modulation of gut ecology in a model industrialized microbiota. The ability to leverage this strain-level adaptation for engraftment provides a framework for understanding functional divergence between non-industrialized and industrialized microbiomes.

Although many gut bacterial species are distributed across geographically and culturally diverse populations^6–9^, the metabolic traits of strains carried by non-industrialized populations remain poorly characterized. A limited number of strain-resolved metagenomic studies have identified population-associated phylogenetic and genomic structure among the *Segatella copri* complex and *Bifidobacterium* spp.^5,8–10,34,35^. Yet linking such genetic variation to metabolic phenotypes remains challenging, in part because cultured isolates from traditional populations, which enable direct functional validation, remain scarce. This gap is particularly important because the accessory gene repertoires, comprising genes present in some but not all strains of a species, can underlie adaptive differences in diverse phenotypes relevant to host biology, including nutrient utilization and metabolic transformations^10,12^. Together, these observations prompted us to investigate within-species functional diversity among prevalent gut bacteria. Our genomic analyses revealed population-associated phylogenetic structure and differences in predicted functional repertoires across multiple Clostridia species shared between Hadza and industrialized populations (Figures 1E and 1G). This phylogenetic structure was not restricted to Hadza-industrialized comparisons; for example, *G. qucibialis* genomes from Asian cohorts also displayed geographic clustering distinct from those of other populations (Figure S1B), consistent with previous reports showing strain diversification across populations^7–9^. Notably, Hadza MAGs from the majority of prevalent Clostridia species examined contained fewer annotated KEGG functions and CAZyme families than their non-Hadza counterparts (Figures 1G and 2C), despite the greater taxonomic diversity previously reported in the Hadza microbiota^4,6^. Hadza MAGs also had shorter assembled genome lengths in 11 of the 17 Clostridia species analyzed in the current study, which may partly account for their reduced functional richness. One possible interpretation is that taxonomic loss during industrialization is accompanied by a redistribution of functional repertoires among retained taxa, potentially allowing them to encode a wider range of community functions^36–38^. Consistent with this possibility, species enriched in industrialized microbiomes, such as those of the *Bacteroides* genus, often encode extensive machinery for utilizing diverse dietary and host-derived glycans^39,40^. As exceptions to this overall trend, Hadza MAGs from several Clostridia species were enriched for specific metabolic functions, including pathways involved in amino acid and carbohydrate metabolism in *D. longicatena*. (Figures 1G and 2A). Functional assays using cultured isolates further demonstrated corresponding population-associated differences in carbohydrate utilization (Figure 3D). These findings therefore reveal an additional layer of functional diversity within the Hadza microbiota that is obscured in conventional community-level metagenomic analyses.

Recent population-genomic studies suggest that gut bacteria adapt to diet and other host-associated factors at the subspecies level. Genome-wide sweeps can generate ecologically distinct lineages, whereas gene-specific sweeps can spread adaptive loci across divergent strain backgrounds through homologous recombination^41,42^. Targets of gene-specific sweeps are enriched for carbohydrate-related genes, including those encoding GHs; these targets also differ between industrialized and non-industrialized populations^42^. Our observations are consistent with this evolutionary framework, revealing both Hadza-enriched genetic variation and distinct carbohydrate-utilization phenotypes in *D. longicatena*. In particular, the localized sequence conservation of the GH43/GH5 PUL between *D. longicatena* and a Hadza-associated VANISH *Dorea* species, together with its mobile-element architecture, is consistent with carbohydrate-utilization machinery being exchanged among related species and maintained within a population, potentially under shared dietary pressures (Figures 4B–4D). Although these observations do not establish a gene-specific selective sweep, they support a model in which population-restricted taxa serve as reservoirs of accessory metabolic functions for species shared across populations.

Additionally, our study expands the diversity and population specificity of CAZymes and putative PULs in gut Clostridia. Studies of gut polysaccharide utilization have historically centered on *Bacteroides* and *Bifidobacterium*, which have become model glycan degraders^43,44^. By contrast, polysaccharide utilization by gut Clostridia has been understudied, with relatively few well-characterized examples, including resistant starch degradation by *Ruminococcus bromii*^45^, β-mannan degradation by *Roseburia intestinalis*^46^, and Gram-positive PULs in *Roseburia* spp. and *Eubacterium rectale*^47^. This knowledge gap is notable because many fiber-intervention studies have identified Clostridia, including major producers of short-chain fatty acids (SCFAs)^48,49^, among the strongest responders^50–53^. These findings emphasize the need to define how Clostridia participate in gut polysaccharide metabolism. We found that many prevalent Clostridia encoded CAZymes predicted to act on plant cell-wall glycans and fructans, but their prevalence shows a different tendency between Hadza and non-Hadza MAGs (Figures 2D and S2A). Plant cell-wall-related CAZymes were enriched in several Hadza-associated Clostridia, whereas fructan-related CAZymes were generally enriched in industrialized Clostridia. This population-dependent pattern is directionally consistent with a recent intervention in which a non-industrialized-type high-fiber diet enriched several GH43 subfamilies involved in plant cell-wall degradation^54^. These findings show that gut Clostridia harbor diverse polysaccharide-degradation repertoires that vary across populations and strains. We further validated utilization of pectic arabinan by Hadza *D. longicatena* isolates (Figures 3D and 3E). In *D. longicatena*, however, arabinan-responsive CAZymes and transporter genes were encoded at separate loci, in contrast to canonical *Bacteroides* PULs, in which CAZymes, transporters, and regulators are co-localized^25^. This organization suggests that *D. longicatena* may release oligosaccharides and monosaccharides extracellularly before uptake, rather than employing the canonical *Bacteroides* “selfish” mechanism in which oligosaccharides are imported for periplasmic degradation^55^.

Taken together, our study reveals a previously unresolved layer of functional diversity within gut bacterial species shared between Hadza and industrialized populations and demonstrates that diet-responsive metabolic machinery from Hadza strains can be applied to modulate gut ecology in a model industrialized microbiota. These findings establish a framework for identifying strain-specific metabolic traits across diverse gut Clostridia and testing their impact on community ecology and host biology, potentially enabling precision microbiome interventions informed by bacterial metabolism.

### Limitation of the Study

First, we focused primarily on *D. longicatena* as a representative species for experimental validation of the population-associated metabolic repertoires predicted from Hadza Clostridia MAGs, in part because many strictly anaerobic Clostridia remain difficult to isolate and culture. *D. longicatena* exhibited one of the strongest population-associated differences in CAZyme repertoires (Figure 2A), making it a strong candidate for experimental investigation; nevertheless, further studies are needed to characterize the metabolic capabilities of other Clostridia species. Second, although Hadza dietary fiber intake has been estimated to be four to eight times that of typical U.S. population^21^, the abundance and structural composition of arabinan in the Hadza diet remain unknown. Therefore, a direct relationship between habitual dietary exposure and enrichment of arabinan-utilization machinery cannot yet be established. Finally, we could not genetically validate the genes underlying arabinan utilization because genetic tools are available for only a few Lachnospiraceae species^56^. Although comparative genomic analyses across multiple isolates identified Hadza-enriched candidate genes associated with these phenotypes, genetic manipulation will be required to establish causality.

## Resource availability

### Lead contact

Further information and requests for resources should be directed to and will be fulfilled by the lead contact, Justin Sonnenburg.

### Materials availability

This study did not generate new unique reagents.

### Data and code availability

- Any additional information required to reanalyze the data reported in this paper is available from the lead contact upon request.

## Acknowledgements

We are indebted to the Hadza as well as the numerous people and organizations who provided support that have enabled this project. We also thank Surya Tripathi, Sean Spencer, Matthew Olm, Sailendharan Sudakaran, Maggie Madrigal-Moeller, and Kate Miller for their helpful advice and support, Michael Fischbach, Mikhail Iakiviak, and the Microbiome Therapies Initiative (MITI) for providing bacterial strains and genome assemblies, Isolation Bio for providing access to its technology and technical support, and BENEO for providing reagents. The metabolome analysis of this study was made possible by the Chan Zuckerberg Biohub. J.L.S. is a Chan Zuckerberg Biohub Investigator. The study was supported by Coefficient Giving, The Algard family, National Institutes of Health grants R01-DK085025, R01-AT013330 (J.L.S.), and 1K99DK146142 (T.T.). The content is solely the responsibility of the authors and does not necessarily represent the official views of the National Institutes of Health.

## Author contributions

Conceptualization, T.T. and J.L.S.; methodology, T.T., A.R., E.K.R, S.K.H, E.D.S., and J.L.S.; software, T.T.; investigation, T.T., A.R., K.K., and E.K.R.; resources, S.K.H., E.D.S. and J.L.S.; writing – original draft, T.T. and J.L.S.; writing – review & editing, T.T., E.D.S., and J.L.S.; visualization, T.T.; funding acquisition, T.T., E.D.S., and J.L.S.; supervision, E.D.S. and J.L.S.; project administration, J.L.S.

## Methods

### Experimental model and study participant details

#### Gnotobiotic mouse models

All animal procedures were approved by the Stanford University Institutional Animal Care and Use Committee and performed in accordance with protocol 19727. The animal facility was maintained at 21–22 °C and 30–80% relative humidity under a 12-h light/12-h dark cycle. Germ-free Swiss Webster and C57BL/6 mice were bred and maintained in vinyl germ-free isolators and fed an autoclaved standard diet (Purina LabDiet 5K67). Mice were matched by sex and age and were 8–12 weeks old at the start of the experiments.

To establish gnotobiotic mouse models, germ-free mice were orally gavaged with 200 µL of a mixture of individual bacterial cultures comprising the 15-member synthetic community (15SM; excluding *D. longicatena*). To facilitate colonization, mice received a second 200 µL gavage containing a mixture of Clostridia species 2–4 days later. After bacterial colonization was confirmed by qPCR, mice were switched to a diet lacking microbiota-accessible carbohydrates (Envigo; Teklad custom diet TD.150689), containing glucose (696 g/kg) as the sole carbohydrate source^14,57^. One week later, polysaccharide supplementation in the drinking water was initiated (see below for the preparation method), and the experimental bacterial strains were administered by oral gavage at a volume of 200 µL per mouse. Mice had ad libitum access to food and drinking water, including polysaccharide-supplemented water during the intervention period. Polysaccharide-supplemented drinking water was replaced twice weekly. Fecal samples were collected two to three times per week until the experimental endpoint.

At necropsy, mice were anesthetized with ketamine hydrochloride (100 mg/kg) and xylazine (5–15 mg/kg) administered via an intraperitoneal injection. Portal venous blood was collected by portal vein puncture into serum-separation tubes and maintained at room temperature for 60 min to allow clot formation. Following blood collection, mice were euthanized by cervical dislocation under deep anesthesia, and intestinal contents were collected. Blood samples were centrifuged at 1,500 g for 10 min to separate the serum. Serum, intestinal-content, and fecal samples were stored at −80 °C.

### Method details

#### Metagenome-assembled genome database and microbiota prevalence analysis

Fecal metagenomic data and metagenome-assembled genomes (MAGs) generated in a previous study of Hadza hunter-gatherers and comparative populations were used^6^. Briefly, ultra-deep shotgun metagenomic sequencing generated 388 metagenomic datasets from 351 fecal samples collected from 167 Hadza individuals, with an average sequencing depth of 24.2 Gbp per metagenomic dataset. For comparative analyses, fecal metagenomes from 56 Nepali individuals and 12 Californian individuals were also included. Quality-filtered reads were assembled using SPAdes v3.13. Bacterial and archaeal genomes were recovered using a co-mapping approach followed by binning with MetaBAT2 v2.15. Genome bins were refined using MAGpurify v2 and retained if they had ≥50% estimated completeness and <10% estimated contamination. The archived dataset comprised 54,779 bacterial and archaeal MAGs.

The bacterial and archaeal genomes were clustered into species-level groups at ≥95% average nucleotide identity over ≥30% of their lengths using dRep v3.0.0, and a species representative genome (SRG) was selected from each cluster. Metagenomic reads were mapped to the SRG database using Bowtie2, and the resulting alignments were processed using inStrain v1.2.14. An SRG was considered detected in a sample when its coverage breadth was ≥0.5. For the present analysis, SRGs detected in ≥50% of samples from a given population were considered prevalent.

Phylogenetic analysis of SRGs was performed using GToTree v1.8.6^58^. GToTree additionally filtered genomes according to the proportion of lineage-specific marker genes detected, as specified by the -G parameter. Genomes were retained if they contained at least 50% of the bacterial marker genes, using the parameters -H Bacteria and -G 0.5. Phylogenetic trees were visualized using the R packages ggtree v4.0.5^59^ and ggplot2 v4.0.3. Singleton SRGs were removed from the analysis. The definitions of VANISH and BloSSUM taxa were described previously^6^. Briefly, VANISH (volatile and/or associated negatively with industrialized societies of humans) and BloSSUM (bloom or selected in societies of urbanization/modernization) taxa were species preferentially detected in Hadza and industrialized populations, respectively. These assignments were based on Fisher’s exact tests comparing species prevalence between the two populations, with species in the ≥95th percentile of ranked significance scores assigned according to the direction of enrichment. The UpSet plot summarizing the prevalence of SRGs (Figure 1B) was made using the R packages ComplexUpset v1.3.3 and ggplot2 v3.5.2. SRGs with a prevalence of at least 10% in either the Hadza or U.S. population were included in the plot.

#### Inclusion criteria and quality control of metagenome-assembled genomes

Genomes were obtained from our previous Hadza study^6^, which generated 54,779 MAGs, and from the Unified Human Gastrointestinal Genome database (UHGG) v2.0.2^23^, which contains 278,263 MAGs and 10,969 isolate genomes. Taxonomic assignment was performed using GTDB-Tk v1.7.0 with GTDB reference database release 202^60^.

To investigate strain-level divergence among prevalent species, we initially identified 32 SRGs that were prevalent in both the Hadza and North America populations. One SRG, Blautia_A wexlerae_A, was subsequently excluded because closer examination indicated that its apparent prevalence resulted from erroneous taxonomic assignments between Blautia_A wexlerae_A and Blautia_A wexlerae, likely owing to their close phylogenetic relationship. Subsequent analyses therefore focused on the remaining 31 SRGs (Figure S1A). For each SRG, genomes from four population groups, i.e., Hadza, North America, Europe, and Asia, were included (Figure 1D). Other populations were excluded because of the limited number of available MAGs. MAGs from U.S. and Nepali participants generated in the Hadza study were assigned to the North America and Asia groups, respectively, whereas the remaining genomes were obtained from the UHGG database.

For downstream analyses, MAGs were subjected to the following quality-control criteria: (1) ≥80% completeness and ≤5% contamination, consistent with previous comparative genomic studies using MAGs^5,61–63^; and (2) at least 20 MAGs from each of the four population groups for a given SRG to provide sufficient group representation. MAG quality was assessed using CheckM2 v1.1.0^64^, and the resulting completeness and contamination estimates were included as covariates in all association models, as described below. After quality control, 17 SRGs that were prevalent in both the Hadza and U.S. populations met the inclusion criteria and were retained for subsequent analyses. The global map and population-composition pie charts (Figure 1D) were generated in R using the R packages rnaturalearth v1.2.0, sf v1.1-1, countrycode v1.8.0, and ggplot2 v4.0.3.

#### Phylogenetic analysis and ANI dendrogram

Phylogenetic analyses were performed using GToTree v1.8.6^58^. For each SRG, all MAGs that passed the quality-control criteria described above were included. Genomes were retained if they contained at least 75% of the Firmicutes-specific marker genes, using the parameters -H Firmicutes and -G 0.75.

During analysis of the UHGG genomes, MGYG000197446, which was annotated as *Dorea*_A *longicatena* in the UHGG database, appeared as a marked phylogenetic outlier relative to other genomes assigned to the same GTDB taxon. BLAST analyses using BLAST v2.12.0 identified 18 contigs with taxonomic assignments inconsistent with D. longicatena. For example, contig_5 (92,436 bp), contig_7 (85,530 bp), and contig_10 (69,259 bp) matched *Coprococcus comes* ATCC 27758 with 100% query coverage and an E-value of 0. In contrast, the best matches to *D. longicatena* genomes had query coverages of only 32%, 0%, and 14%, respectively. The 18 putatively contaminating or mis-binned contigs accounted for 25.9% of the total assembly length. After removal of these contigs, the estimated genome quality was 61.4% completeness and 0.0% contamination. Because the resulting genome no longer met our predefined inclusion criterion of ≥80% completeness, MGYG000197446 was excluded from subsequent phylogenetic and comparative genomic analyses, leaving 16,959 genomes for downstream analysis.

Pairwise average nucleotide identity (ANI) among *Dorea* genomes was calculated using dRep v3.4.5^65^ with FastANI v1.34^66^ as the secondary comparison algorithm. Primary clustering was performed using Mash with a sketch size of 100,000 and a primary ANI threshold of 0.95. FastANI was run with its default parameters. ANI distance was calculated as 1 − ANI, with ANI expressed as a proportion, and the resulting distance matrix was visualized using ggplot2 v4.0.3.

Between-population phylogenetic distances were quantified from species-specific phylogenetic trees generated using GToTree. For each species, pairwise patristic distances among genome tips were calculated from branch lengths using the cophenetic.phylo function in the R package ape v5.8-1. Genomes were assigned to four population groups, i.e., Hadza, North America, Europe, and Asia, and the distance between each pair of populations was calculated as the mean of all pairwise distances between genomes from the two populations. Mean distances for the six population pairs were standardized within each species and visualized as a heatmap using pheatmap v1.0.13. Species and population pairs were hierarchically clustered using Euclidean distance and average linkage. To identify species showing greater Hadza-associated phylogenetic separation, the six population-pair mean distances for each species were analyzed using an exploratory weighted linear regression comparing the three population pairs that included the Hadza with the three pairs that did not. Each population-pair mean was weighted by the harmonic mean of the corresponding population sample sizes. Positive regression coefficients indicated greater mean phylogenetic distances in population pairs that included the Hadza. P values were adjusted across species using the Benjamini–Hochberg method, and adjusted P < 0.05 was considered significant.

#### KEGG module completeness analysis

MAGs assigned to species detected across Hadza and industrialized populations were analyzed for KEGG functional content. Protein-coding sequences were predicted from MAGs using Prodigal v2.6.3^67^. To reduce redundancy before functional annotation, predicted proteins from all MAGs were clustered with MMseqs2 v17-b804f at 95% amino-acid identity and 80% alignment coverage (--min-seq-id 0.95 -c 0.8 --cov-mode 1)^68^. Representative sequences were annotated with KofamScan v1.3.0^69^, and KEGG ortholog (KO) assignments were propagated to the corresponding cluster members. Within each MAG, multiple occurrences of the same KO were collapsed to binary presence. MAGs were filtered based on the quality and prevalence criteria described above.

KEGG module completeness was calculated using the R package mclink v1.1.2^70^. KEGG module definitions were evaluated according to their logical structure: comma-separated alternative KOs were treated as OR relationships, whereas plus-separated required KOs were treated as AND relationships. KEGG module definitions and Level 3 classifications were obtained from the KO_pathway_ref dataset distributed with mclink.

#### CAZyme annotation and substrate classification

CAZymes were annotated using dbCAN v4.1.4 in a containerized environment (tadtake/dbcan4_nf) with dbCAN HMMdb v12 and the dbCAN-sub HMM database released in 2022^71,72^. Genome assemblies were provided as input for MAGs, whereas predicted protein FASTA files were used for bacterial isolate genomes. For MAG-based comparative analyses, we used CAZyme annotations identified by HMMER against dbCAN HMMdb. Domain-coordinate information appended to the HMMER annotations was removed, and each unique HMMER assignment, including multidomain combinations, was treated as a distinct CAZyme type. We identified 510 distinct CAZyme types across all included MAGs from four populations, with a mean of 34.7 types per MAG (SD, 15.7). Types detected in only one MAG across the combined dataset were excluded from downstream profile and type-specific comparative analyses.

For functional comparisons, distinct CAZyme types were defined from cleaned dbCAN HMMER assignments and counted once per MAG, irrespective of protein copy number. CAZyme types containing glycoside hydrolases (GHs), polysaccharide lyases (PLs), or carbohydrate esterases (CE) domains were classified as predominantly degradative (degradation-type), whereas those containing glycosyltransferases (GT) but no GH/PL/CE domain were classified as predominantly biosynthetic (biosynthesis-type). Auxiliary activity (AA) enzymes and carbohydrate-binding modules (CBMs) contributed to total CAZyme type richness but were not used to assign types to these functional categories.

Predicted CAZyme genes were assigned to substrate classes using a manually curated CAZyme–substrate list compiled from published annotations and additional literature review^73^. Pectin/hemicellulose- and cellulose-associated glycans were combined into a single plant cell-wall glycan category, and human milk oligosaccharide (HMO) was combined into a host glycan category, resulting in six analyzed substrate classes: plant cell-wall glycans, fructan, starch-associated glycans, fungal glycans, host glycans, and algal glycans. Substrate assignment was restricted to GH, PL, and CE domains to focus the analysis on predicted polysaccharide-degrading activities. Exact family or subfamily annotations were used when available; otherwise, substrate annotations were inherited from the corresponding parent family. CAZyme types associated with more than one broad substrate class were considered ambiguous and excluded from substrate-specific analyses. Each gene was counted once and assigned only when all informative GH, PL, and CE domains indicated the same broad substrate class. Genes containing ambiguous or conflicting GH, PL, or CE annotations were excluded. Putative substrate classes were assigned to 61% and 51% of the GH/PL/CE-containing CAZyme types evaluated in the MAG and metagenomic analyses, respectively.

#### Metagenomic read mapping to a non-redundant gene catalog

Protein-coding sequences were predicted from the Hadza MAG collection and genomes included from the UHGG v2.0.2 using Prodigal v2.6.3. Genome identifiers were added to the predicted coding-sequence headers before concatenation. Sequences shorter than 100 bp were removed using SeqKit v2.10.0.

Non-redundant gene catalogs were initially generated separately for the Hadza and UHGG genome collections using MMseqs2 v17.b804f easy-linclust with a minimum nucleotide sequence identity of 95% and a coverage threshold of 90% (--min-seq-id 0.95 --cov-mode 1 -c 0.9 --cluster-mode 2). Because of the size of the UHGG dataset, subsets of UHGG genes were first clustered independently, after which their representative sequences were pooled and reclustered using the same parameters. Representative sequences from the Hadza and UHGG catalogs were then combined and subjected to a final round of clustering using the same parameters. The resulting non-redundant catalog contained 20,920,795 representative gene sequences, with a total sequence length of approximately 19.1 Gbp.

Quality-controlled paired-end metagenomic reads from adult Hadza, Nepali, and U.S. participants were subsampled before mapping to reduce differences in sequencing depth. The rarefaction depth was selected as the highest one-million-read-pair increment that retained at least 90% of eligible samples from each population, resulting in a target of approximately 44 million read pairs per sample. Reads were randomly subsampled with SeqKit using a sample-specific proportion calculated from the original number of unique read pairs. The same random seed was applied to both mates to preserve read pairing. The resulting libraries contained approximately 41.7–44.0 million read pairs.

Subsampled paired-end reads were mapped and quantified against the nonredundant gene catalog using Salmon v1.10.3^74^ through a custom Nextflow workflow implemented in Nextflow v24.10.4. A Salmon index was generated from the representative nucleotide sequences using a k-mer length of 31. Gene-level abundance was quantified using automatic library-type detection (-l A), mapping validation (-- validateMappings), and metagenomic mode (--meta). Mapping rates were calculated as the percentage of processed fragments mapped to the catalog. Four samples with mapping rates below 70% were excluded. After these exclusions, 317 metagenomes were retained, comprising 249 Hadza, 56 Nepali, and 12 U.S. samples, with population-specific mean mapping rates ranging from 80.2% to 84.9%. Following catalog-wide quantification, Salmon estimated fragment counts (NumReads) for reference genes with dbCAN HMMER annotations were retained for downstream CAZyme analyses.

#### Bacterial isolation from human fecal samples

Bacterial strains were obtained through manual and automated isolation. Manual isolation from Hadza fecal samples was performed as described previously^6^. Briefly, fecal suspensions were plated anaerobically on various solid media, including YCFA, MRS, Columbia, and BHI agar. The previous study reported 52 isolates recovered from 14 Hadza fecal samples. These included one isolate that was classified as *Dorea longicatena*_B and designated ST02 in the current study. Further manual isolation expanded the collection to 103 isolates from 19 Hadza fecal samples, including two additional *D. longicatena* isolates. Additionally, 634 bacterial isolates were recovered from two Hadza fecal samples, one from an infant and one from an adult, using the Prospector automated isolation platform at General Automation Lab Technologies (GALT; now Isolation Bio). Prospector enables the parallel cultivation of individual bacterial cells at the nanoliter scale and the transfer of selected clonal isolates into 96-well plates. A total of 413, 192, and 29 isolates were recovered using YCFA-C, BHI, and GAM media, respectively. Isolates were taxonomically identified using the MALDI Biotyper system (Bruker) and Sanger sequencing of 16S rRNA gene amplicons. Based on the MALDI Biotyper and 16S rRNA gene annotations, eight isolates were provisionally classified as *D. longicatena* and selected for whole-genome sequencing. Genome-based classification identified seven *D. longicatena* isolates and one *Dorea*_A sp934294775 isolate. Among the *D. longicatena* isolates, Ga09, Ga10, and Ga12 were recovered from the same donor and were nearly identical phylogenetically, with pairwise average nucleotide identities of ≥99.99%. Because these isolates likely represented the same lineage, Ga10 and Ga12 were excluded from subsequent analyses to reduce redundancy. Two additional *D. longicatena* strains were isolated from fecal samples collected from two adult donors in the U.S. and were included as isolates from an industrialized population.

#### Whole genome sequencing

Bacterial strains were streaked from glycerol stocks onto YCFA-C (GMExpression; GMNB-YCFA02) agar plates and cultured overnight under anaerobic conditions. A single colony was picked and subcultured in YCFA-C medium overnight. The bacterial cultures were centrifuged at 6,000 g for 5 min, and the resulting pellets were resuspended in TE buffer. Genomic DNA was extracted using the MasterPure Gram Positive DNA Purification Kit (Biosearch Technologies; catalog no. MGP04100) according to the manufacturer’s instructions. Briefly, bacterial cells were lysed using Ready-Lyse Lysozyme, Proteinase K, and Gram Positive Cell Lysis Solution. Proteins and cellular debris were removed using MPC Protein Precipitation Reagent, and residual RNA was digested with RNase A. DNA was subsequently precipitated with isopropanol, washed with 70% ethanol, and resuspended in TE buffer.

Whole-genome sequencing and hybrid genome assembly were performed by Plasmidsaurus, a commercial sequencing service. Briefly, Oxford Nanopore Technologies long-read sequencing and genome assembly were performed, followed by polishing with 150-bp paired-end Illumina short reads using Polypolish v0.6. Taxonomic assignment was performed using GTDB-Tk v2.3.0 with GTDB reference database release 214^60,75^. ANI dendrogram was made as described above. For genomic analysis, VE303_06 and i26_0019_D6, the only complete *D. longicatena* genome assemblies available from NCBI besides the type strain DSM 13814, were also included as representatives of isolates from industrialized populations. Gene prediction was performed using Prodigal v2.6.3. Carbohydrate-active enzymes were annotated using dbCAN v4.1.4 with the dbCAN-sub HMM database, as described above.

#### Pangenome and functional enrichment analysis

A pangenome was constructed using Panaroo v1.5.2^76^ in strict-cleaning and strict gene-refinding modes. Prokka-annotated GenBank files were used as the input files. Initial gene clustering used sequence-identity and length-difference cutoffs of 0.98, whereas family-level clustering used sequence-identity and length-difference cutoffs of 0.70. Core gene families, defined by a prevalence threshold of 1.0, were aligned using MAFFT v7.526^77^. Panaroo was also run without sequence alignment to obtain unaligned nucleotide and protein sequences for subsequent sequence-identity analyses.

Pangenome presence-absence analyses were restricted to ten *D. longicatena* genomes: five Hadza isolates (ST1L, ST02, Ga09, Ga16, and 1E6) and five isolates from industrialized populations (DSM 13814, MITI2, MITI3, VE303_06, and i26_0019_D6). Ga11 was excluded from these summaries because it was assigned to *Dorea*_A sp934294775 rather than *D. longicatena*. Gene families present in all ten *D. longicatena* genomes were classified as core. Non-core families present in at least two Hadza genomes and absent from all industrialized genomes were classified as Hadza-associated accessory families, whereas those present in at least two industrialized genomes and absent from all Hadza genomes were classified as industrialized-associated accessory families. All remaining non-core families were classified as other accessory families. The resulting binary presence-absence matrix was visualized by hierarchical clustering using Euclidean distance and the Ward.D2 linkage method using the R package pheatmap v1.0.13.

Protein-coding genes were assigned to COG functional categories using COGclassifier v2.0.0, and CAZymes were identified using the HMMER results from dbCAN v4.1.4. Gene-level annotations were collapsed to the Panaroo-family level; a COG category was assigned using the most frequent annotation among family members, and a family was classified as CAZyme-encoding when at least one member contained a dbCAN HMM hit. Enrichment of COG categories and CAZyme-encoding families within each pangenome category was evaluated using two-sided Fisher’s exact tests comparing the target category with the remaining background. Tests were conducted using the entire pangenome as the background. Fold enrichment was calculated as the proportion of families carrying the annotation within the target category divided by the corresponding proportion in the background. P values were adjusted within each background using the Benjamini–Hochberg method.

#### Growth curve assay

Depleted medium, a carbohydrate-defined medium based on a previously described YCFA formulation^18^, was used to assess carbohydrate utilization by *D. longicatena*. In addition to the components of *Bacteroides* minimal medium, depleted medium contains yeast extract and beef extract, enabling the growth of certain Clostridia species with amino acid auxotrophies, including *D. longicatena*. For *A. hadrus*, YCFA carbohydrate-defined medium, in which glucose, maltose, and cellobiose were replaced with a single defined carbohydrate, was used because this species did not grow in depleted medium.

Unless otherwise indicated, carbohydrates were added at a final concentration of 0.5% w/v. The following carbohydrates were used: D-glucose (Alfa Aesar; A16828), L-arabinose (Alfa Aesar; A11921), D-fructose (Alfa Aesar; A17718), D-galactose(Acros; 150611000), D-xylose (Sigma-Aldrich; X1500), D-galacturonic acid monohydrate (Sigma-Aldrich; 48280), D-cellobiose (Thermo; 108460250), sucrose (Sigma-Aldrich; S0389), arabinan (Megazyme; P-ARAB), rhamnogalacturonan (Megazyme; P-RHAGA), arabinogalactan (Sigma-Aldrich; 10830), galactan (Megazyme; P-GALPOT), Pectin (Acros; 416862500), Pectin from apple (Sigma-Aldrich; 76282), inulin from dahlia tubers (Sigma-Aldrich; I3754), galactomannan (Megazyme; P-GALML), wheat arabinoxylan (Megazyme; P-WAXYL), xyloglucan (Megazyme; P-XYGLN), and type II mucin (Sigma-Aldrich; M2378). The fructooligosaccharide (FOS) preparations, with a degree of polymerization of 2 to 8, were provided by BENEO-Orafti NV.

Bacterial strains were streaked from glycerol stocks onto YCFA-C agar plates and cultured overnight under anaerobic conditions. A single colony was picked and subcultured overnight in YCFA-C medium. Carbohydrate-defined media were dispensed into 96-well plates at 200 µL per well, and bacterial cultures were inoculated at a 1:100 dilution. Plates were incubated anaerobically at 37 °C for up to 48 h in an Epoch 2 microplate spectrophotometer (BioTek) housed inside a Coy anaerobic chamber. OD600 was measured every 15 min following 30 s of shaking. Media and assay plates were pre-reduced in the anaerobic chamber for at least 24 h before use. Growth data were analyzed using the R package gcplyr v1.12.0^78^.

#### Comparative analysis of gene clusters

Genomic regions of interest were extracted from Prokka-generated GenBank files, with region boundaries defined by the first and last specified locus tags and without additional flanking coding sequences. Homologous gene clusters were identified using cblaster v1.3.19^79^ in local-search mode against DIAMOND databases generated from the annotated genomes. The search of the ST1L GH43/GH146 and GH43/Gh5 loci required a minimum amino acid identity of 50%, with other search and cluster-detection parameters left at their default values. Detected clusters were extracted as GenBank files and compared using clinker v0.0.29^80^, which performed pairwise protein-sequence alignments and visualized gene order, orientation, and sequence identity.

#### Bacterial RNA-sequencing

*Dorea longicatena* ST1L was streaked from a glycerol stock onto a YCFA-C agar plate and cultured overnight under anaerobic conditions. A single colony was picked and subcultured overnight in depleted medium containing 0.5% glucose (w/v). The bacterial culture was centrifuged at 2,200 g for 5 min and resuspended in depleted medium without carbohydrates. The suspension was inoculated at a 1:100 dilution into 2.5 mL of experimental medium consisting of depleted medium supplemented with 0.5% glucose, L-arabinose, or arabinan (w/v) in a 24-well deep-well plate. After 9 h, the cultures were centrifuged at 4,700 g for 5 min, and the pellets were resuspended in PM1 buffer supplemented with 2-mercaptoethanol, according to the instructions provided with the RNA extraction kit described below. The mean final OD600 values were 0.51, 0.33, and 0.23 for the glucose, L-arabinose, and arabinan conditions, respectively.

Total RNA was extracted using the RNeasy PowerMicrobiome Kit (Qiagen; #26000-50) according to the manufacturer’s instructions. The extracted RNA was sent to SeqCenter for RNA sequencing. Briefly, the RNA samples were first treated with RNase-free DNase (Invitrogen). Libraries were prepared using the Illumina Stranded Total RNA Prep Ligation with Ribo-Zero Plus kit for ribosomal RNA depletion and 10-bp unique dual indexes. Sequencing was performed on an Illumina NovaSeq X Plus platform to generate 150-bp paired-end reads. Demultiplexing, quality control, and adapter trimming were performed using bcl-convert v4.2.4. Sequencing generated an average of 2.0 × 10⁷ reads and 2.5 × 10⁹ bases with Phred quality scores >30 per sample.

The demultiplexed paired-end RNA-sequencing reads were further processed using fastp v1.3.6^81^ with default parameters to remove residual adapter sequences and low-quality reads. The genome assembly of *D. longicatena* strain ST1L was used as the reference genome. An index of the ST1L reference genome was generated using BWA-MEM2 v2.3. Quality-filtered paired-end reads were aligned to the indexed reference genome using bwa-mem2 mem. The resulting SAM files were converted to BAM format and coordinate-sorted using SAMtools v1.23.1. For visualization of genome-wide RNA-seq coverage, the sorted BAM files were indexed using SAMtools and converted to bedGraph files using the bamCoverage tool in deepTools v3.5.4^82^. Coverage values were normalized as counts per million mapped reads (CPM).

#### Arabinan hydrolysis by resting *D. longicatena* cells

The capacity of *D. longicatena* ST1L to hydrolyze extracellular arabinan was assessed using washed resting cells and the p-hydroxybenzoic acid hydrazide (PAHBAH) reducing-sugar assay. ST1L and DSM 13814 were inoculated at a 1:200 dilution into 30 mL of YCFA medium supplemented with 0.5% arabinan or glucose (w/v) and cultured anaerobically at 37 °C for 6 h, by which point the cultures had reached an OD600 of approximately 0.4 to 0.6, except ST1L cultured in glucose-containing medium, which achieved an OD600 of approximately 1.0 due to faster growth. Cells were collected by centrifugation at 6,000 g for 5 min and washed twice with pre-reduced PBS to remove residual carbohydrates from the culture medium. The washed cell pellet was resuspended to an OD600 of 5 in pre-reduced PBS containing 0.5% arabinan (w/v) or in PBS alone. Arabinan-containing PBS without cells and PBS alone were included as substrate and buffer controls, respectively. Reactions were incubated anaerobically at 37 °C. Samples were collected at 0, 1, 4, 8, and 24 h after reaction setup.

At each time point, samples were centrifuged at 16,000 g for 5 min to remove bacterial cells and insoluble arabinan. The clarified supernatants were analyzed using the PAHBAH assay^83^. Briefly, 30 µL of each sample, control, or L-arabinose standard was mixed with 90 µL of freshly prepared PAHBAH working reagent and heated at 95 °C for 6 min. Samples were cooled to room temperature for 10 min, and absorbance was measured at 410 nm. A standard curve containing 0–1.0 mM L-arabinose was used to express reducing-sugar release as L-arabinose equivalents. Signals were corrected using matched arabinan-only and buffer-only controls.

#### BLASTp analysis of PUL homologs in *Dorea* MAGs

Protein homologs of the *D. longicatena* ST1L GH43/GH146 and the *Dorea*_A sp934294775 Ga11 GH43/GH5 loci were identified using protein BLAST (BLASTp) implemented in BLAST v2.12.0. Protein sequences were extracted from the Prokka-annotated ST1L and Ga11 protein FASTA files using SeqKit v2.10.0 and queried against a BLAST protein database comprising concatenated Prokka-predicted proteins from Hadza and UHGG Dorea MAGs with ≥90% completeness and ≤5% contamination. BLASTp searches were performed with an E-value threshold of 1 × 10^−5^ and a maximum of 10,000 target sequences per query. Alignments with ≥30% amino acid identity, ≥30% query coverage per high-scoring segment pair, and an E value ≤1 × 10^−5^ were retained. For each query protein and MAG, the highest-scoring alignment was selected according to bit score, followed by query coverage and amino acid identity. Subject coverage was calculated as the alignment length divided by the subject-protein length. Hits with an E value ≤1 × 10−5, ≥40% amino acid identity, and ≥50% coverage of both the query and subject proteins were considered evidence of homolog presence in downstream prevalence analyses.

Population-level prevalence was evaluated for the region spanning ST1L_wgs_02244 to ST1L_wgs_02260 in MAGs assigned to *Dorea*_A *longicatena* and for the region spanning Ga11_wgs_01249 to Ga11_wgs_01264 in MAGs assigned to *Dorea*_A *longicatena* or *Dorea*_A *longicatena*_B. MAG metadata and short Prokka locus-tag identifiers were integrated to determine the population origin of each genome. For the ST1L analysis, MAGs were grouped into Hadza, North America, Europe, Asia, and Oceania populations, whereas the Ga11 analysis compared Hadza and pooled non-Hadza MAGs. Homolog prevalence was calculated as the proportion of MAGs in each population containing a qualifying hit. In the ST1L analysis, prevalence in Hadza MAGs was compared separately with that in each non-Hadza population using two-sided Fisher’s exact tests. In the Ga11 analysis, Hadza and non-Hadza prevalence was compared separately within each species. Comparisons required at least three MAGs in each group, and P values were adjusted across query genes within each population comparison using the Benjamini–Hochberg procedure.

#### Pairwise nucleotide identity analysis of homologous genes

To assess the sequence identity of candidate PULs among Hadza-associated *Dorea* genomes, a pangenome constructed from Panaroo v.1.5.2 with unaligned sequences was used (--clean-mode strict, --refind-mode strict, --len_dif_percent 0.98, --family_len_dif_percent 0.70, -a pan, and --aligner none). The resulting gene_presence_absence.csv and gene_data.csv files were used to identify homologous gene families shared between representative genome pairs, including *D. longicatena* ST1L and *Dorea*_A sp934294775 Ga11. Gene families lacking a gene identifier in either genome or containing multiple semicolon-delimited gene identifiers were excluded, restricting pairwise sequence comparisons to shared gene families represented by a single unambiguous gene in each genome.

For each shared gene family, the corresponding nucleotide and amino acid sequences were retrieved from gene_data.csv. Nucleotide and protein sequences were globally aligned using pairwiseAlignment() from the R package pwalign v1.6.0, and percent sequence identity was calculated using pid() with the PID1 definition. Candidate PULs were defined based on their genomic organization and CAZyme annotations, and the corresponding Panaroo gene families were used to designate PUL-associated genes. Sequence identities of PUL-associated genes were compared with those of shared gene families outside the corresponding PUL.

To determine whether candidate PULs were more highly conserved than expected from genome-wide sequence similarity, empirical permutation tests were performed separately for nucleotide and protein identity. For each PUL gene set, an equal number of non-PUL gene families was randomly sampled without replacement from the genomic background, and the median identity, mean identity, and proportion of genes with ≥95% sequence identity were calculated. This procedure was repeated 100,000 times. Empirical one-sided P values were calculated as (k+1)/(N+1), where k was the number of permutations yielding a statistic greater than or equal to the observed value and N was the total number of permutations. Nucleotide sequence identity was visualized according to genomic position using ggplot2 v4.0.3.

#### IS-like elements flanking the GH43/GH5 PUL

Analysis of the ST1L genomic region surrounding the GH43/GH5 PUL identified two oppositely oriented 1,308-bp genes annotated by Prokka as IS4-family transposases (ST1L_wgs_01881 and ST1L_wgs_01906). For each gene, the transposase and 3 kb of flanking sequence on either side were extracted using SeqKit v2.10.0, generating two 7,308-bp regions. The region containing ST1L_wgs_01906 was reverse-complemented to orient the two sequences consistently. The regions were compared by global alignment using Needle from EMBOSS v6.6.0^84^ and by local alignment using BLASTn from BLAST v2.12.0, which identified a highly conserved region of approximately 1.7 kb encompassing each transposase and approximately 200 bp of flanking sequence on either side.

Terminal inverted repeats (TIRs) were evaluated in extended 2,301-bp regions using einverted from EMBOSS v6.6.0 and independently using Generic Repeat Finder (GRF) v1.0.2 in TIR-detection mode. GRF searches used terminal-repeat lengths of 10–80 bp, spacer lengths of 1,400–1,750 bp, a 10-bp seed permitting one mismatch, and maximum allowances of 20 mismatches and five indels. GRF identified a 1,691-bp candidate IS-like element surrounding each transposase, together with associated TIRs. The two IS-like elements shared 99.4% nucleotide identity. TIR sequences defined from the GRF-predicted boundaries were globally aligned using the R package pwalign, and complementarity was calculated as the proportion of complementary nucleotide pairs after excluding alignment columns containing indels.

#### VANISH *Dorea* gene-family sharing across *Dorea* species

To define protein families in the Hadza-associated VANISH species *Dorea*_A sp934294775, Prokka-predicted proteins from 25 high-quality MAGs (≥90% completeness and ≤5% contamination) were clustered using MMseqs2 v17-b804f easy-cluster, with ≥70% amino acid identity, ≥80% bidirectional coverage, set-cover clustering (--cluster-mode 0), and cluster reassignment. Families present in ≥90% of source MAGs were classified as core, whereas those present in two or more but <90% were classified as accessory; singletons were excluded.

Family representatives were searched using BLASTp v2.12.0 against Prokka-predicted proteins from high-quality *Dorea*_A *longicatena*, *Dorea*_A *longicatena*_B, and *Dorea formicigenerans* MAGs, using an E-value threshold of 1 × 10^−10^, one high-scoring segment pair per subject, and up to 10,000 target sequences per query. For each family and recipient MAG, the best alignment was selected based on hit class, bit score, query coverage, subject coverage, and amino acid identity. Hits with ≥40% identity and ≥70% coverage of both proteins were considered homologs. For each recipient MAG, the proportions of VANISH core and accessory families detected were calculated. Hadza and non-Hadza MAGs were compared using separate linear models for each recipient species and family group, adjusting for assembled genome length, completeness, and contamination. Adjusted population means were obtained using estimated marginal means, and P values were corrected across recipient species within each family group using the Benjamini–Hochberg method.

Family representatives were further annotated using dbCAN v4.1.4 with dbCAN HMMdb v12. Positive HMMER assignments were grouped as glycan-degradation-related CAZymes (GH, PL, and CE) or glycan-biosynthesis-related CAZymes (GT); AA and CBM families were excluded. CAZyme-family sharing was analyzed using the same homolog criteria and covariate-adjusted linear models. Hadza–non-Hadza differences were obtained from estimated marginal means, with Benjamini–Hochberg correction across CAZyme categories within each recipient species and family group.

#### Preparation of sugar beet pulp extract for animal studies

To prepare crude arabinan solution for animal studies, 375 g of sugar beet pulp (SBP; Premium Beet Pulp Shreds; Standlee Premium Products) was suspended in 1.5 L of water and homogenized in two batches using a food processor for approximately 5 min. An additional 1.5 L of water was then added to obtain a total volume of 3 L. The suspension was autoclaved for 40 min at 121°C and cooled to room temperature. The autoclaved suspension was passed through cheesecloth to remove large insoluble particles. The filtrate was clarified by centrifugation twice at 7,000 g for 10 min. The clarified supernatant was typically 1.4 L, depending on evaporation and clarification. It was diluted to 2.0 L before sterilization by passage through a 0.22-µm membrane filter.

The arabinan-equivalent content of the SBP extract was estimated by trifluoroacetic acid (TFA) hydrolysis followed by quantification of released L-arabinose^85^. Briefly, 100 µL of SBP extract was mixed with an equal volume of 4 M TFA to obtain a final TFA concentration of 2 M. Samples were heated in sealed glass vials at 120 °C for 90 min and allowed to cool completely. TFA was removed by evaporation under a nitrogen stream at approximately 37 °C. The dried hydrolysates were resuspended in 100 µL of ultrapure water and dried again under nitrogen to remove residual TFA, followed by reconstitution in 500 µL of ultrapure water. L-arabinose released by hydrolysis was quantified using an enzymatic L-arabinose/D-galactose assay (Megazyme; K-ARGA) according to the manufacturer’s instructions. A water-plus-TFA blank and an L-arabinose recovery control were processed through the complete hydrolysis and evaporation procedure. Free L-arabinose was measured in an unhydrolyzed aliquot of the SBP extract at the same effective dilution and subtracted from the total L-arabinose measured after hydrolysis. The resulting bound L-arabinose content was converted to arabinan-equivalent content using a factor of 0.88, which accounts for the mass difference between free arabinose and anhydro-arabinose residues within the polymer. Results were expressed as arabinan-equivalent mass per unit volume of SBP extract. Following TFA hydrolysis, the SBP extract contained 1.9 mg/mL arabinan-equivalent, whereas free L-arabinose in the unhydrolyzed extract was below the assay detection limit.

#### Bacterial strains for gnotobiotic models

The 14-member synthetic community (14SM), together with *D. longicatena* (collectively, 15SM), comprises gut bacterial species commonly found in industrialized populations, including the U.S., and was previously used to model the effects of diet on gut microbial community structure^30,31,86^. Strains were streaked from glycerol stocks onto YCFA-C agar plates and cultured anaerobically at 37 °C for 48–96 h in a Coy anaerobic chamber. Single colonies were picked and subcultured in YCFA-C medium for 48 h. Individual cultures were combined in equal volumes without normalization for cell density because several Clostridia strains did not reach an OD600 greater than 0.2. This approach is consistent with previous gnotobiotic studies in which an established defined bacterial community was used to evaluate microbiota responses to subsequent dietary perturbations^87^. All mice within each experiment received the same community inoculum, which was pooled immediately before oral administration, and the experimental strains and polysaccharides were administered only after establishment of the background community. Therefore, unequal initial abundances of individual community members were shared across treatment groups and were not expected to systematically confound comparisons of the subsequent interventions on *D. longicatena*.

#### *In vitro* community experiments

Depleted medium supplemented with 1% carbohydrate (w/v) was used as the experimental medium, and 15SM was used as the background community for the in vitro experiments. Individual 15SM strains were prepared as described above for the in vivo experiments. *Dorea longicatena* ST1L and DSM 13814 were cultured similarly in YCFA-C medium. Individual cultures were combined in equal volumes without normalization for cell density. The experimental medium was dispensed into 96-well deep-well plates at 800 µL per well, and the bacterial mixtures were inoculated at a 1:200 dilution. Plates were incubated anaerobically at 37 °C for 48 h in an incubator housed within a Coy anaerobic chamber. The medium and assay plates were pre-reduced in the anaerobic chamber for at least 24 h before use. After 48 h of incubation, the final OD600 was measured, and the plates were centrifuged at 6,000 g for 5 min to collect bacterial pellets. The pellets were resuspended in TE buffer and stored at −20 °C until DNA extraction.

#### Fecal microbiota composition analysis of defined bacterial communities

Fecal DNA was extracted using a previously described enzymatic digestion protocol with minor modifications. Briefly, fecal samples were homogenized in TE buffer and sequentially treated with lysozyme (Roche; 10837059001; 300 mg/mL) and a mixture of SDS (Promega; V6551) and proteinase K (Sigma P2308; 25 mg/mL). Samples were incubated for 60 min at 37 °C following the addition of lysozyme and for 30 min at 55 °C following the addition of SDS and proteinase K. DNA was extracted and purified using phenol:chloroform alcohol (UltraPure; Thermo Fisher Scientific; 15593031) and precipitated using 3 M sodium acetate (Sigma S7899) and 100% ethanol. The resulting DNA pellet was dissolved in 100 µL of TE buffer.

The relative abundances of members of the defined bacterial communities were quantified by qPCR using species-specific primers. Primers for members of the 15SM were designed to target species-specific genes or genomic regions present as a single copy per genome, as determined through whole-genome alignments using Large-Scale Genome Alignment Tool (LASTZ) available with Geneious Prime v2025.0 (Geneious). For species shared with the previously described 14-member synthetic community, published primer sets were used. For species-level detection of *D. longicatena*, primers targeting the xanthine phosphoribosyltransferase gene, which is conserved in *D. longicatena* ST1L and DSM 13814, were used to quantify both strains. For strain-specific detection, primers targeting ST1L_01978, a hypothetical gene, and DSM 13814_01159, a hypothetical gene, respectively, were used to quantify each strain within the same community. Cross-reactivity was evaluated for all primer-species combinations by qPCR using DNA extracted from individual bacterial cultures. qPCR was performed using the CFX Connect Real-Time System (Bio-Rad) with Brilliant III Ultra-Fast SYBR Master Mix (Agilent; 600883). Standard curves were used to estimate gene copy numbers from qPCR Ct values. The relative abundance of each bacterial strain was calculated by dividing its estimated copy number by the sum of the copy numbers of all community members detected in the same sample. The absolute abundances of ST1L and DSM 13814 were determined using standard curves generated from Ct values of DNA extracted from bacterial cultures with known CFU counts on YCFA-C agar.

#### Statistical analysis

Comparisons between two groups were performed using Welch’s t-tests. Count data were compared between two groups using two-sided exact Wilcoxon rank-sum tests implemented in the R package coin v1.4-3. Comparisons involving more than two groups were performed using one-way analysis of variance (ANOVA) or two-way factorial ANOVA, followed by the indicated post hoc multiple-comparison test. Associations between categorical variables in contingency tables were evaluated using two-sided Fisher’s exact tests. Where multiple hypotheses were tested, P values were adjusted using either the Benjamini–Hochberg procedure to control the false discovery rate (FDR). Unless otherwise stated, P < 0.05 or adjusted P < 0.05 was considered statistically significant.

#### KEGG module and ortholog analysis

KEGG Level 3 module completeness was analyzed by quasi-binomial logistic regression, including population, MAG completeness, and contamination as predictors. For each genome, completeness was averaged across modules assigned to the same KEGG Level 3 functional category, with equal weighting of modules. Category-species combinations were retained when all four populations were represented, module completeness had at least two distinct values, at least five observations differed from the modal value, and at least ten observations each were above zero and below one. Estimated completeness and contamination were included as covariates to account for residual variation in MAG quality^88^. Hadza was compared with a composite non-Hadza estimate in which North America, Europe, and Asia were weighted according to their numbers of MAGs for each species. Adjusted Level 3 module completeness was predicted at the median MAG completeness and contamination, and effect sizes were reported as Hadza-minus-non-Hadza differences in percentage points. P values were adjusted using the Benjamini–Hochberg procedure across all category-species combinations, separately for the weighted contrast and each population-specific comparison. Associations with adjusted P < 0.05 were considered significant. The 15 significant Level 3 categories associated with the largest numbers of species were displayed, with ties ranked by median absolute effect size (Figure 1G).

Targeted KO-level analyses were performed for tryptophan biosynthesis (M00023) and D-galacturonate degradation (M00631). KO presence was analyzed for species represented by at least 20 MAGs per population and for KOs present and absent in at least five MAGs each. Bias-reduced binomial logistic regression implemented in the R package brglm2 v1.1.0 was used, with population, MAG completeness, and contamination as predictors. Adjusted KO prevalence was predicted at median MAG completeness and contamination. P values from two-sided Wald tests were adjusted using the Benjamini–Hochberg procedure separately within each targeted module and population comparison. A species-KO association was designated consistently enriched or depleted in Hadza only when all three population-specific comparisons had adjusted P < 0.05 and concordant effect directions.

#### CAZyme profiles in metagenome-assembled genomes

All statistical analyses were performed in R v4.5.3 using the R packages MASS v7.3-65, brglm2 v1.1.0, and emmeans v2.0.3. Analyses were performed separately within each species using genomes from the Hadza, North America, Europe, and Asia populations. MAG completeness and contamination estimates were included as covariates in all regression models to account for differences in genome quality.

To analyze population-centroid analysis of CAZyme profiles, binary CAZyme profiles (i.e., presence or absence of CAZymes within each MAG) were averaged within each population for each species to generate prevalence centroids. Bray-Curtis distances were calculated between the four population centroids without additional binary transformation using the R package vegan v2.7-5. Hadza-specific separation was quantified as the mean of the three Hadza–non-Hadza centroid distances minus the mean of the three non-Hadza distances. Significance was evaluated for each species by permuting population labels among MAGs 9999 times while preserving population sample sizes and recalculating the centroids and separation statistic. Two-sided permutation P values were calculated relative to the median null statistic and adjusted across species using the Benjamini–Hochberg method.

CAZyme type richness was analyzed separately within each species using quasi-Poisson regression with a log link, with population, MAG completeness, and contamination as predictors. Hadza estimates were compared with the sample-size-weighted mean of adjusted estimates for three non-Hadza populations. P values were adjusted across species within each architecture panel using the Benjamini–Hochberg method. Type-specific presence was analyzed using bias-reduced logistic regression with the same covariates. Point size represents the fraction of tested CAZyme types whose significant enrichment or depletion agreed with the direction of the corresponding richness difference.

Species-specific counts of putative polysaccharide-degradation CAZyme genes assigned to each substrate class were analyzed using quasi-Poisson regression with a log link, whereas the relative fraction assigned to each substrate class was analyzed using quasi-binomial regression with a logit link and all substrate-predicted GH-, PL-, and CE-containing proteins as the denominator. Each predicted gene contributed one count regardless of the number of domains it contained, and multiple genes with the same CAZyme architecture within a MAG were retained as separate copies. Population, MAG completeness, and contamination were included as predictors in both models. Before modeling, species–substrate combinations were retained only when all four populations contained at least 20 eligible MAGs and the response had at least two observed levels with at least five nonmodal observations. Relative-fraction analyses additionally required at least five MAGs with nonzero focal-substrate counts and at least five with nonzero complementary-substrate counts. Model-adjusted Hadza estimates were compared with the sample-size-weighted mean estimates for the three non-Hadza populations, and P values were adjusted across tests using the Benjamini–Hochberg method. Aggregate plant-polysaccharide-associated counts comprised proteins assigned to plant cell-wall glycans, fructan, or starch-associated glycans.

CAZyme types were defined by the complete dbCAN HMMER annotations assigned to individual genes, including both single-domain and multidomain annotations, as described above. Species-specific copy numbers of individual CAZyme types were analyzed using quasi-Poisson regression with a log link. Each predicted CAZyme-encoding gene contributed one count, and multiple genes of the same type within a MAG were retained as separate copies. Population, MAG completeness, and contamination were included as predictors. Species–CAZyme type combinations were retained only when all four populations contained at least 20 eligible MAGs, the response had at least two observed levels with at least five nonmodal observations, and at least five MAGs had nonzero counts. Model-adjusted Hadza estimates were compared with the sample-size-weighted mean across the three non-Hadza populations, and P values were adjusted across tests using the Benjamini–Hochberg method. CAZyme types showing significant differences in the same direction in at least three species were retained for visualization. Putative substrate classes were assigned when all informative GH, PL, and CE domains within the complete annotation indicated the same broad substrate class, as described above.

#### CAZyme analysis of metagenome-mapped reads

For overall CAZyme richness, a dbCAN-annotated reference gene was considered detected when its Salmon estimated fragment count was at least one. Each detected reference gene was counted once, regardless of the number of HMM profiles assigned to it. CAZyme types were defined by complete dbCAN HMM profile annotations, including family and subfamily profiles and all other profiles present in the HMMER output, such as AA and CBM homology domains. For each type, estimated fragment counts were summed across all reference genes assigned to that profile, and the type was considered detected when the aggregate count was at least one. Types not detected in any retained metagenome were excluded, whereas types detected in a single metagenome were retained.

Population differences in detected reference-gene richness and CAZyme type richness were analyzed using quasi-Poisson regression with a log link. An omnibus F test was used to evaluate the population term, followed by pairwise comparisons among the three populations. Pairwise P values were adjusted separately for each richness measure using the Benjamini–Hochberg method. To assess concordance in population-associated CAZyme differences, each CAZyme type was analyzed separately for detection in Hadza versus Nepali and Hadza versus U.S. metagenomes using quasi-binomial regression with a logit link. Effect estimates were expressed as log_2_ odds ratios for detection in Hadza relative to each non-Hadza population. CAZyme types tested in both comparisons were retained, and concordance between the two sets of effect estimates was evaluated using Pearson’s correlation.

For substrate composition analysis, catalytic annotations were restricted to putative polysaccharide-degradation CAZymes: GH, PL, and CE families. Exact CAZyme family or subfamily substrate annotations were used when available. Types without exact annotations inherited the substrate annotations of their parent families. Substrates were grouped into plant cell-wall, fructan, starch, fungal glycan, host glycan, and algal glycan classes. CAZyme types assigned to more than one of these final broad substrate classes were excluded from the substrate composition analysis. Substrate representation was compared separately between Hadza and Nepali metagenomes and between Hadza and U.S. metagenomes using quasi-binomial regression with a logit link. A substrate class was tested only when all three populations contained at least five samples, the response had at least two observed levels and five nonmodal observations, and every population contained at least five samples with nonzero focal-substrate counts and at least five with nonzero complementary-substrate counts. Effects are reported as log_2_ odds ratios for Hadza relative to the comparison population, with 95% confidence intervals. P values were adjusted across substrate classes separately within each population comparison using the Benjamini–Hochberg method.

#### Longitudinal data analysis

A repeated-measures ANOVA was performed for longitudinal data, including competitive index and abundance in animals, using the R packages afex v1.5.-1 and emmeans v2.0.4. Briefly, competitive index and abundance values were log_10_-transformed before statistical analysis. The ANOVA models included strain and carbohydrate conditions as between-subject factors and day as a within-subject factor. Biological replicate identity was included as the subject variable to account for repeated measurements over time. Type III sums of squares were used, and the Greenhouse-Geisser correction was applied to effects involving day to account for potential violations of sphericity. Significant interactions were followed by simple-effects analyses based on estimated marginal means, comparing carbohydrate conditions within each strain at each day. All tests were two-sided, and Holm-adjusted P < 0.05 was considered statistically significant.

