## Supplemental Figures for "Functional Divergence in Gut Clostridia from Hadza Hunter-Gatherers Reveals Diet-Adapted Metabolic Machinery That Shapes Gut Ecology"

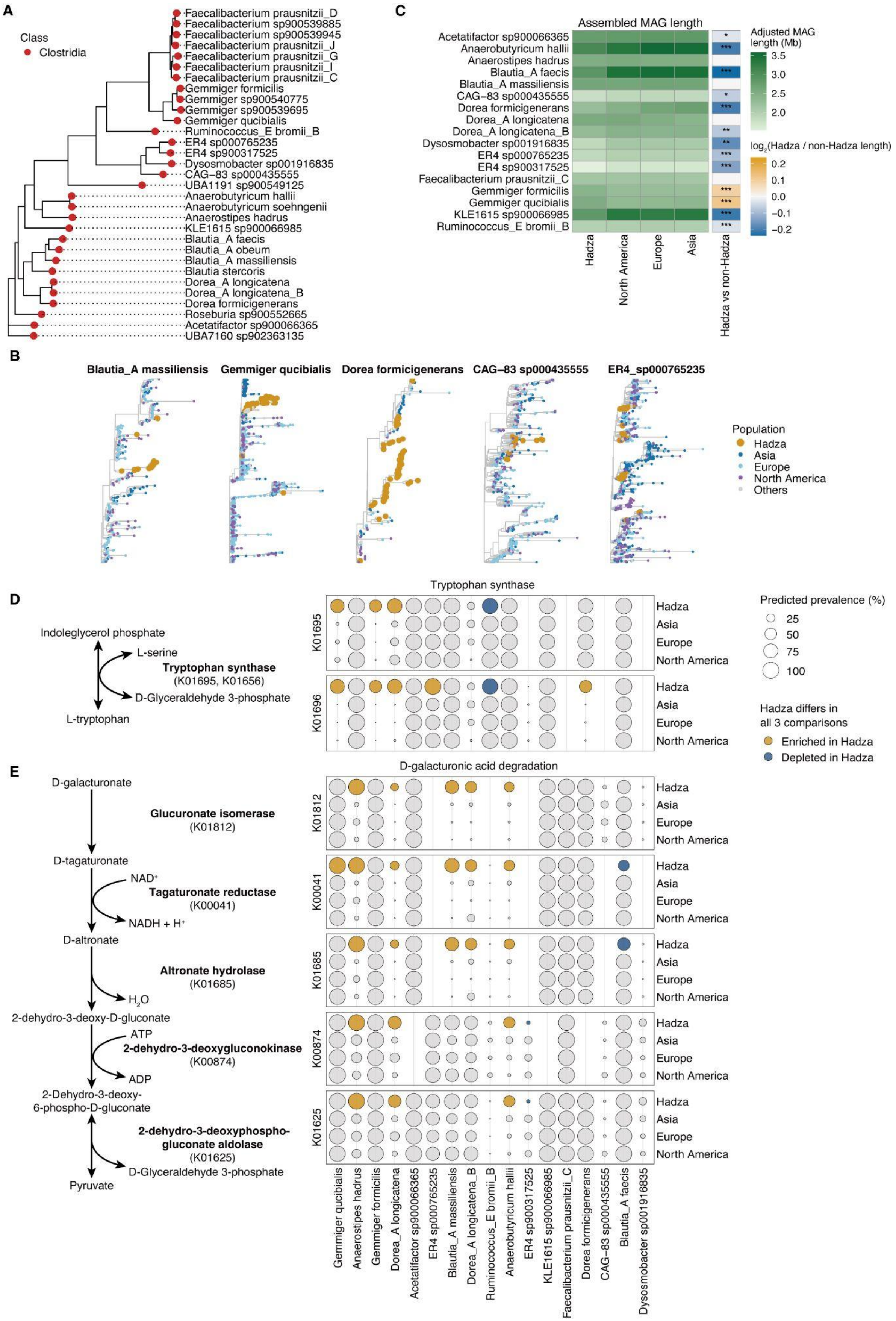
A
Class
Clostridia
Faecalibacterium prausnitzii_D
Faecalibacterium sp900539885
Faecalibacterium sp900539945
Faecalibacterium prausnitzii_J
Faecalibacterium prausnitzii_G
Faecalibacterium prausnitzii_I
Faecalibacterium prausnitzii_C
Gemmiger formicilis
Gemmiger sp900540775
Gemmiger sp900539695
Gemmiger qucibialis
Ruminococcus_E bromii_B
ER4 sp000765235
ER4 sp900317525
Dysosmobacter sp001916835
CAG−83 sp000435555
UBA1191 sp900549125
Anaerobutyricum hallii
Anaerobutyricum soehngenii
Anaerostipes hadrus
KLE1615 sp900066985
Blautia_A faecis
Blautia_A obeum
Blautia_A massiliensis
Blautia stercoris
Dorea_A longicatena
Dorea_A longicatena_B
Dorea formicigenerans
Roseburia sp900552665
Acetatifactor sp900066365
UBA7160 sp902363135
C
Assembled MAG length
Acetatifactor sp900066365
Anaerobutyricum hallii
Anaerostipes hadrus
Blautia_A faecis
Blautia_A massiliensis
CAG−83 sp000435555
Dorea formicigenerans
Dorea_A longicatena
Dorea_A longicatena_B
Dysosmobacter sp001916835
ER4 sp000765235
ER4 sp900317525
Faecalibacterium prausnitzii_C
Gemmiger formicilis
Gemmiger qucibialis
KLE1615 sp900066985
Ruminococcus_E bromii_B
Hadza
North America
Europe
Asia
Hadza vs non-Hadza
Adjusted MAG length (Mb)
3.5
3.0
2.5
2.0
log2(Hadza / non-Hadza length)
0.2
0.1
0.0
−0.1
−0.2
B
Blautia_A massiliensis
Gemmiger qucibialis
Dorea formicigenerans
CAG−83 sp000435555
ER4_sp000765235
Population
Hadza
Asia
Europe
North America
Others
D
Indoleglycerol phosphate
L-serine
Tryptophan synthase
(K01695, K01656)
D-Glyceraldehyde 3-phosphate
L-tryptophan
Tryptophan synthase
K01695
K01696
Hadza
Asia
Europe
North America
Predicted prevalence (%)
25
50
75
100
Hadza differs in all 3 comparisons
Enriched in Hadza
Depleted in Hadza
E
D-galacturonate
Glucuronate isomerase
(K01812)
D-tagaturonate
NAD+
Tagaturonate reductase
(K00041)
NADH + H+
D-altronate
Altronate hydrolase
(K01685)
H2O
2-dehydro-3-deoxy-D-gluconate
ATP
2-dehydro-3-deoxygluconokinase
(K00874)
ADP
2-Dehydro-3-deoxy-6-phospho-D-gluconate
2-dehydro-3-deoxyphospho-gluconate aldolase
(K01625)
D-Glyceraldehyde 3-phosphate
Pyruvate
D-galacturonic acid degradation
K01812
K00041
K01685
K00874
K01625
Gemmiger qucibialis
Anaerostipes hadrus
Gemmiger formicilis
Dorea_A longicatena
Acetatifactor sp900066365
ER4 sp000765235
Blautia_A massiliensis
Dorea_A longicatena_B
Ruminococcus_E bromii_B
Anaerobutyricum hallii
ER4 sp900317525
KLE1615 sp900066985
Faecalibacterium prausnitzii_C
Dorea formicigenerans
CAG−83 sp000435555
Blautia_A faecis
Dysosmobacter sp001916835

**Figure S1. Phylogenetic and Functional Uniqueness of Hadza Clostridia MAGs (Related to Figure 1)**
(A) Marker-gene-based phylogenetic tree showing the 31 Clostridia species prevalent in both Hadza and U.S. populations, as shown in Figure 1A.
(B) Marker-gene-based phylogenetic trees of Clostridia species exhibiting significant Hadza-biased divergence in Figure 1E, including *Blautia*_A *massiliensis*, *Gemmiger qucibialis*, *Dorea formicigenerans*, CAG−83 sp000435555, and ER4_sp000765235. Colors of dots show the origins of MAGs. Dot colors indicate the population origins of the MAGs.
(C) Heatmap showing population-associated differences in assembled MAG length. Mean MAG length adjusted for completeness and contamination is shown. The sidebar shows the $\log_2$ ratio between Hadza and sample-size-weighted non-Hadza estimates.
(D and E) Metabolic pathway schematics (left) and population-level predicted prevalence (right) of KEGG orthologs constituting tryptophan synthase (D) and the D-galacturonic acid degradation pathway (E) in Clostridia MAGs. Hadza bubbles are colored when KO prevalence was significantly higher (blue) or lower (red) in Hadza in all three population-specific comparisons with concordant effect directions.
Linear regression (C) and bias-reduced binomial logistic regression (D and E) were used for statistical analysis. *adjusted $P < 0.05$; **adjusted $P < 0.01$; ***adjusted $P < 0.001$.

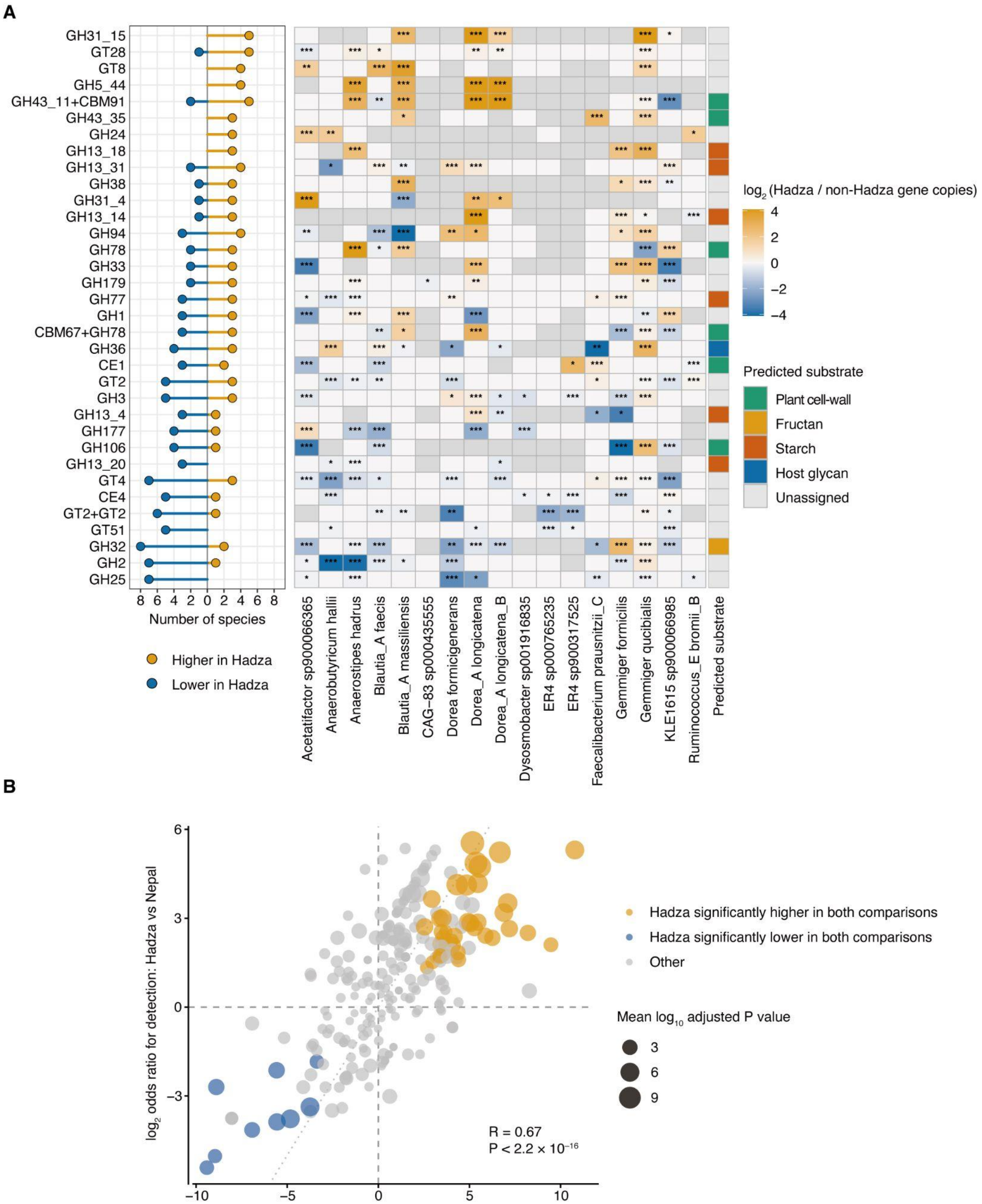


**Figure S2. CAZyme Diversity and Uniqueness in Hadza Clostridia MAGs (Related to Figure 2)**

(A) Population-associated differences in CAZyme gene copy numbers per MAG within each species. The lollipop plot (left) shows the number of species in which the copy number of each CAZyme family was significantly higher or lower in Hadza MAGs. The heatmap (right) shows species-specific $\log_2$ ratios of adjusted counts in Hadza MAGs to the weighted non-Hadza reference. Orange and blue indicate higher and lower counts in Hadza MAGs, respectively; white indicates tested but nonsignificant comparisons, and gray indicates comparisons not tested because too few MAGs were retained for statistical analysis. The sidebar indicates putative substrate classes.

(B) Concordance of population-associated differences in CAZyme type detection. Each point represents a CAZyme type. Colored points indicate types differing significantly in the same direction in both comparisons. The dotted diagonal indicates the line of identity, along which the estimated $\log_2$ odds ratios are equal for the two comparisons. The result of Pearson's correlation is also described.

Quasi-Poisson regression with a log link (A) and bias-reduced logistic regression (B) were used. *adjusted $P < 0.05$; **adjusted $P < 0.01$; ***adjusted $P < 0.001$.

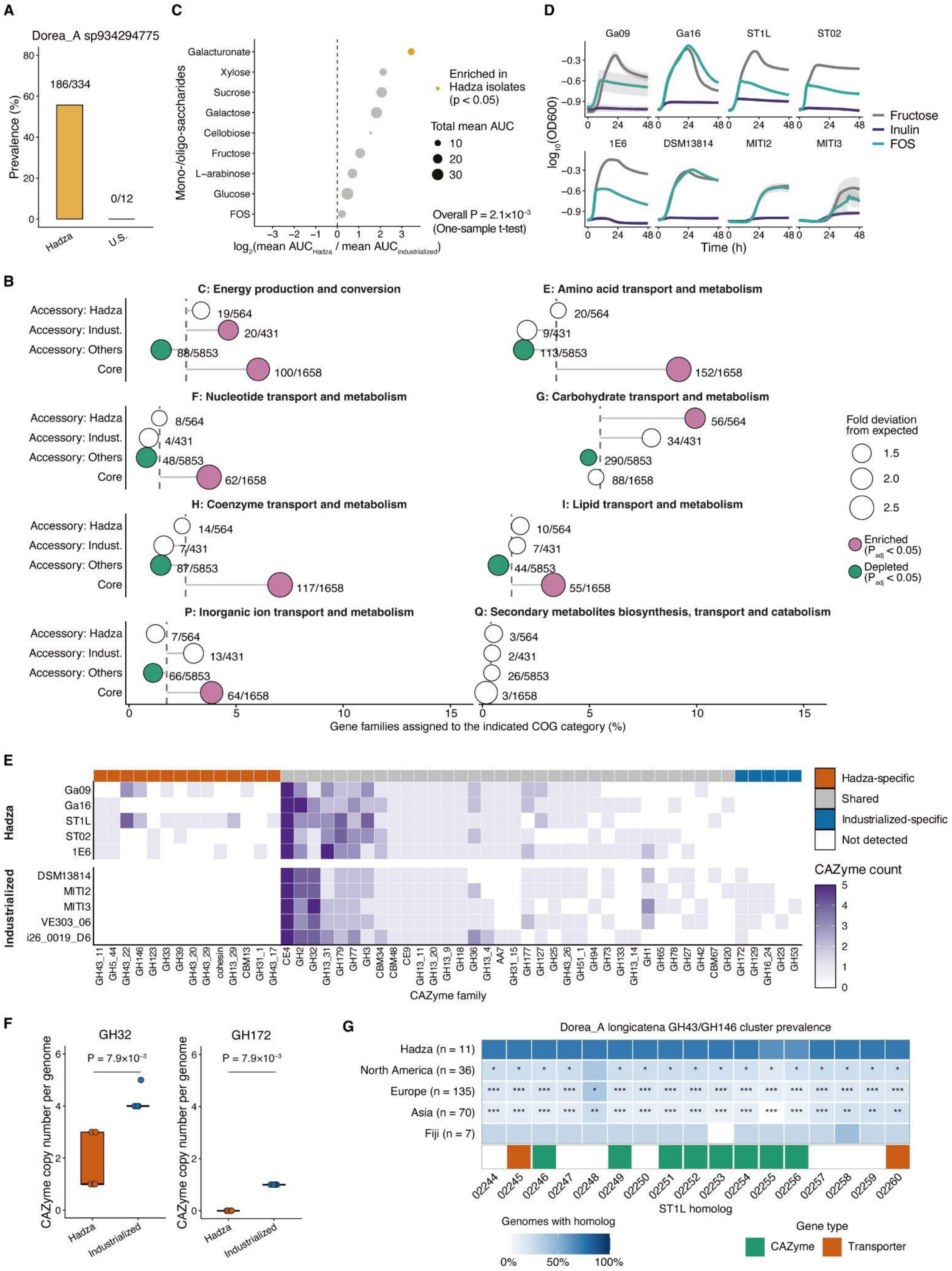
A
Dorea_A sp934294775
Prevalence (%)
186/334
0/12
Hadza
U.S.
C
Mono-/oligo-saccharides
Galacturonate
Xylose
Sucrose
Galactose
Cellobiose
Fructose
L−arabinose
Glucose
FOS
Enriched in Hadza isolates (p < 0.05)
Total mean AUC
10
20
30
Overall P = 2.1×10⁻³ (One-sample t-test)
log2(mean AUCHadza / mean AUCindustrialized)
D
Ga09
Ga16
ST1L
ST02
1E6
DSM13814
MITI2
MITI3
log10(OD600)
Time (h)
Fructose
Inulin
FOS
B
C: Energy production and conversion
E: Amino acid transport and metabolism
F: Nucleotide transport and metabolism
G: Carbohydrate transport and metabolism
H: Coenzyme transport and metabolism
I: Lipid transport and metabolism
P: Inorganic ion transport and metabolism
Q: Secondary metabolites biosynthesis, transport and catabolism
Accessory: Hadza
Accessory: Indust.
Accessory: Others
Core
19/564
20/431
88/5853
100/1658
20/564
9/431
113/5853
152/1658
8/564
4/431
48/5853
62/1658
56/564
34/431
290/5853
88/1658
14/564
7/431
87/5853
117/1658
10/564
7/431
44/5853
55/1658
7/564
13/431
66/5853
64/1658
3/564
2/431
26/5853
3/1658
Gene families assigned to the indicated COG category (%)
Fold deviation from expected
1.5
2.0
2.5
Enriched (Padj < 0.05)
Depleted (Padj < 0.05)
E
Hadza
Ga09
Ga16
ST1L
ST02
1E6
Industrialized
DSM13814
MITI2
MITI3
VE303_06
i26_0019_D6
CAZyme family
Hadza-specific
Shared
Industrialized-specific
Not detected
CAZyme count
F
GH32
GH172
P = 7.9×10⁻³
P = 7.9×10⁻³
CAZyme copy number per genome
Hadza
Industrialized
G
Dorea_A longicatena GH43/GH146 cluster prevalence
Hadza (n = 11)
North America (n = 36)
Europe (n = 135)
Asia (n = 70)
Fiji (n = 7)
ST1L homolog
Genomes with homolog
0%
50%
100%
Gene type
CAZyme
Transporter

**Figure S3. Carbohydrate Utilization and CAZyme Diversity in Hadza D. longicatena Isolates (Related to Figure 3)**

(A) Prevalence of *Dorea*_A sp934294775 in the Hadza and U.S. populations.

(B) Proportions of COG-annotated gene families among four gene-prevalence categories. COG categories related to metabolism are shown. The proportion in each category was compared with the genome-wide expected proportion. Significant enrichment and depletion relative to the expected proportion are indicated in pink and green, respectively.

(C) Growth potential of *D. longicatena* isolates cultured with mono- and oligosaccharides, expressed as the ratio of the mean AUC of Hadza isolates to that of industrialized isolates. Disk size indicates the overall mean AUC, and disk color indicates significantly greater growth among Hadza isolates.

(D) Growth curves of eight *D. longicatena* isolates cultured in depleted medium supplemented with 0.5% fructose, inulin, or fructooligosaccharide (FOS). Shaded areas indicate the standard deviation (n = 3 per group).

(E) Copy numbers of CAZymes except biosynthesis-type in ten *D. longicatena* isolate genomes, including two complete genome assemblies from NCBI. The top bar indicates the population specificity of each CAZyme, defined by its absence from all isolates from the opposite population.

(F) GH32 and GH172 copy numbers in ten *D. longicatena* isolate genomes.

(G) Local sequence conservation of homologs of the GH43/GH146 cluster from Hadza *D. longicatena* strain ST1L in *D. longicatena* genomes from North America, Europe, Asia, and Fiji. The white-to-blue color scale indicates homolog prevalence. Asterisks denote significant differences between Hadza and non-Hadza populations.

Fisher's exact test (B and G), one-sample t-test of the $\log_2$-transformed ratios against zero (C), and two-sided exact Wilcoxon rank-sum test (F) were used for statistical analysis. *adjusted $P < 0.05$; **adjusted $P < 0.01$; ***adjusted $P < 0.001$.

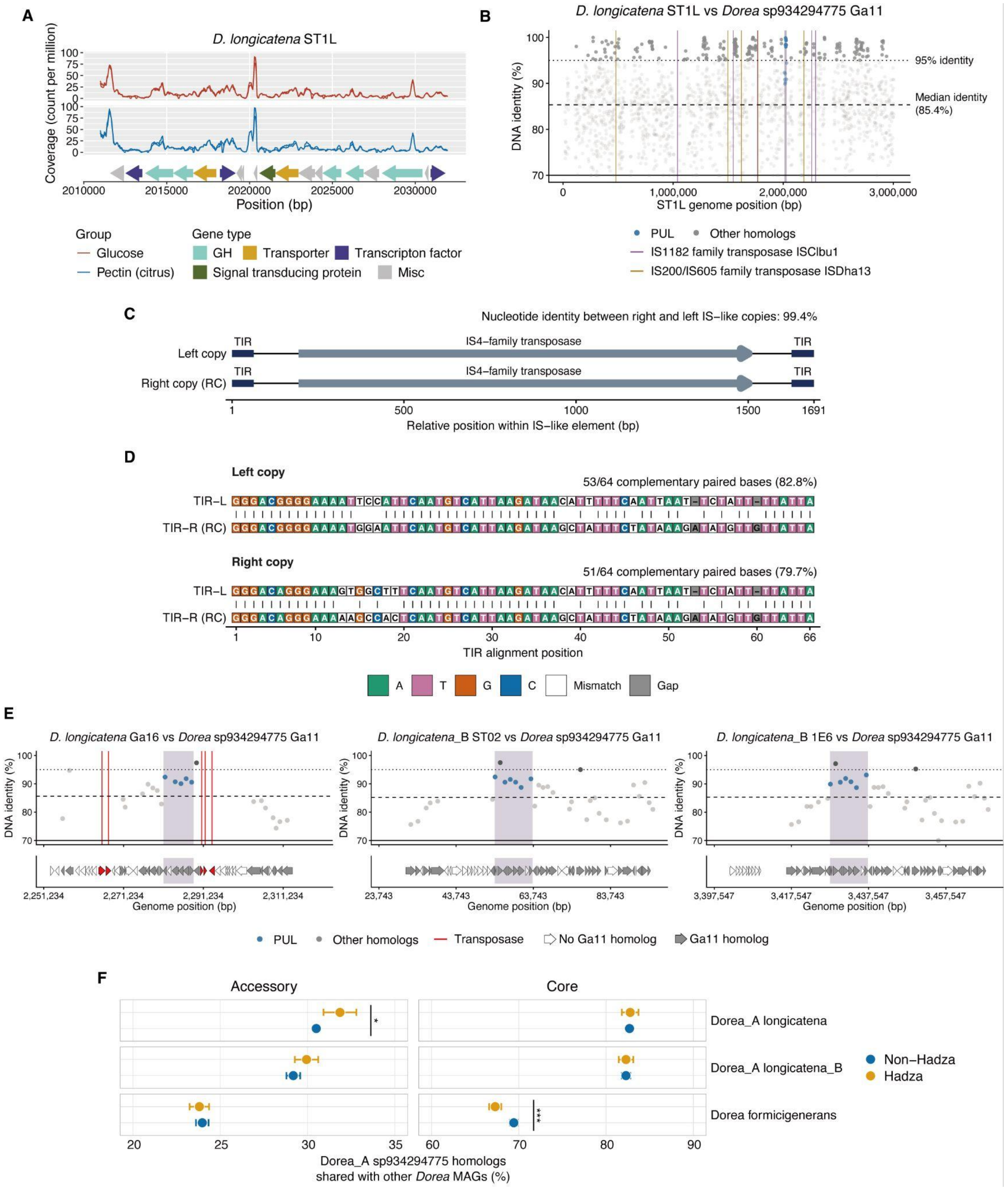


**Figure S4. Sequence Conservation and Mobile-Element Structure Associated with a PUL in Hadza *D. longicatena* Isolates (Related to Figure 4)**

(A) RNA-sequencing analysis showing the transcriptional activity of the GH43/GH5 PUL in Hadza *D. longicatena* strain ST1L. Depleted medium supplemented with 1% of the indicated carbohydrate was used (n = 2 per group).

(B) Genome-wide map of Hadza *D. longicatena* strain ST1L showing the nucleotide identities of homologs shared with Hadza *Dorea*_A sp934294775 strain Ga11. Purple and orange lines indicate the locations of the IS1182-family transposase ISClbu1 and the IS200/IS605-family transposase ISDha13, respectively. Blue dots indicate homologs from the GH43/GH5 PUL.

(C) Schematic showing the structures and homology of two IS-like copies flanking the GH43/GH5 PUL region from Hadza *D. longicatena* strain ST1L. The IS-like copies comprise IS4-family transposases and their corresponding imperfect terminal inverted repeats (TIRs).

(D) Alignment and sequence identity of the TIRs flanking the two IS4-family transposases.

(E) Magnified genome maps of Hadza *D. longicatena* strain Ga16, *D. longicatena*_B strain ST02, and *D. longicatena*_B strain 1E6, showing the nucleotide identities of homologs shared with Hadza *Dorea*_A sp934294775 strain Ga11. Red lines indicate the locations of transposase gene families, and blue dots indicate homologs from the GH43/GH5 PUL. Gray and white gene arrows indicate homologs shared with Ga11.

(F) Sharing of core and accessory homologs between Hadza *Dorea*_A sp934294775 genomes and Hadza or industrialized genomes from three *Dorea* species. Core homologs are defined as those with ≥90% prevalence among Hadza Dorea_A sp934294775 genomes. Accessory homologs are defined as non-core, non-singleton homologs. Linear regression was used for statistical analysis (F). *adjusted $P < 0.05$; ***adjusted $P < 0.001$.

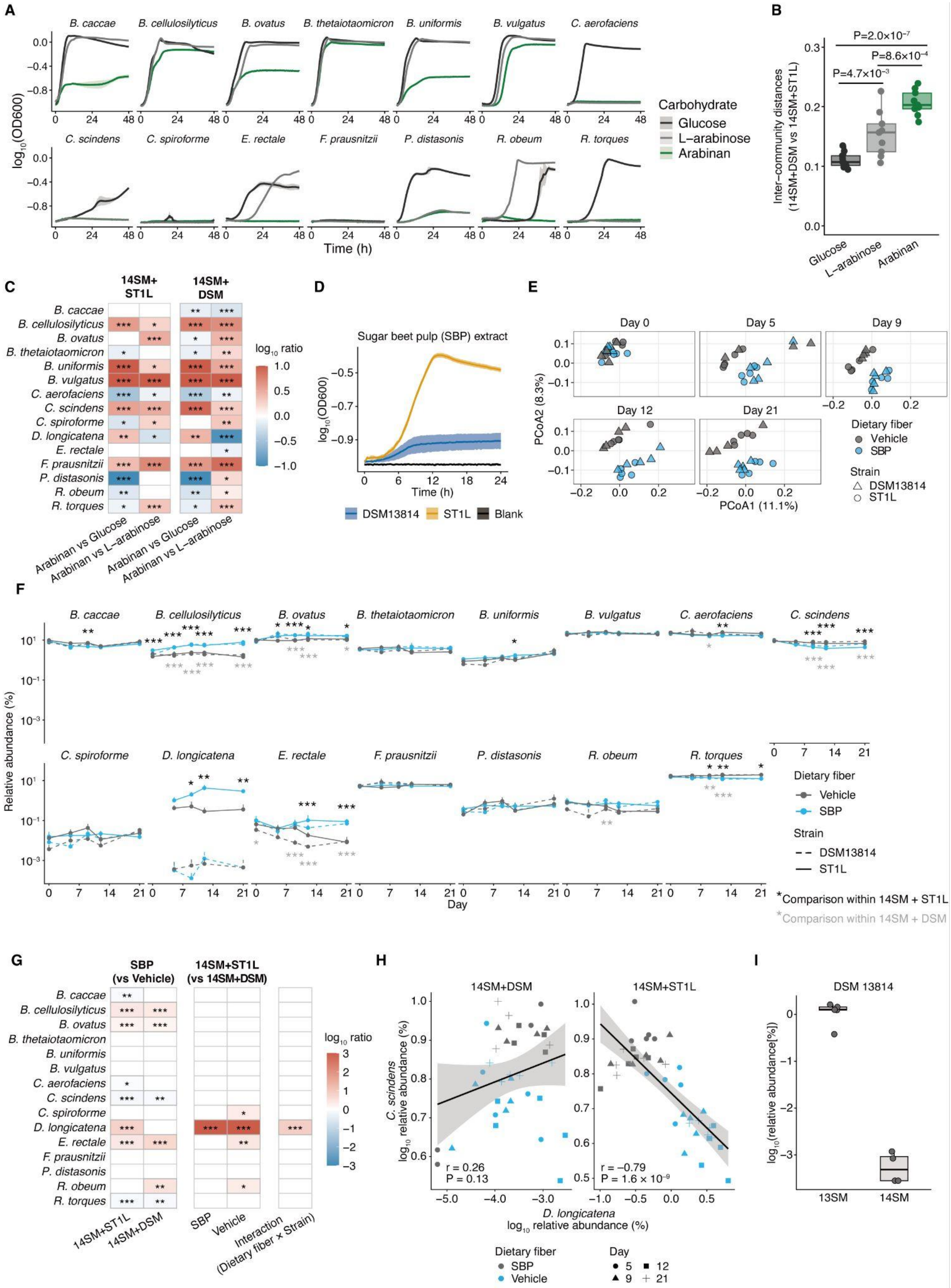
A
B. caccae
B. cellulosilyticus
B. ovatus
B. thetaiotaomicron
B. uniformis
B. vulgatus
C. aerofaciens
C. scindens
C. spiroforme
E. rectale
F. prausnitzii
P. distasonis
R. obeum
R. torques
log10(OD600)
Time (h)
Carbohydrate
Glucose
L–arabinose
Arabinan
B
P=2.0×10−7
P=8.6×10−4
P=4.7×10−3
Inter–community distances (14SM+DSM vs 14SM+ST1L)
Glucose
L–arabinose
Arabinan
C
14SM+ ST1L
14SM+ DSM
B. caccae
B. cellulosilyticus
B. ovatus
B. thetaiotaomicron
B. uniformis
B. vulgatus
C. aerofaciens
C. scindens
C. spiroforme
D. longicatena
E. rectale
F. prausnitzii
P. distasonis
R. obeum
R. torques
log10 ratio
Arabinan vs Glucose
Arabinan vs L–arabinose
Arabinan vs Glucose
Arabinan vs L–arabinose
D
Sugar beet pulp (SBP) extract
log10(OD600)
Time (h)
DSM13814
ST1L
Blank
E
Day 0
Day 5
Day 9
Day 12
Day 21
PCoA2 (8.3%)
PCoA1 (11.1%)
Dietary fiber
Vehicle
SBP
Strain
DSM13814
ST1L
F
B. caccae
B. cellulosilyticus
B. ovatus
B. thetaiotaomicron
B. uniformis
B. vulgatus
C. aerofaciens
C. scindens
C. spiroforme
D. longicatena
E. rectale
F. prausnitzii
P. distasonis
R. obeum
R. torques
Relative abundance (%)
Day
Dietary fiber
Vehicle
SBP
Strain
DSM13814
ST1L
*Comparison within 14SM + ST1L
*Comparison within 14SM + DSM
G
SBP (vs Vehicle)
14SM+ST1L (vs 14SM+DSM)
B. caccae
B. cellulosilyticus
B. ovatus
B. thetaiotaomicron
B. uniformis
B. vulgatus
C. aerofaciens
C. scindens
C. spiroforme
D. longicatena
E. rectale
F. prausnitzii
P. distasonis
R. obeum
R. torques
log10 ratio
14SM+ST1L
14SM+DSM
SBP
Vehicle
Interaction (Dietary fiber x Strain)
H
14SM+DSM
14SM+ST1L
C. scindens log10 relative abundance (%)
r = 0.26
P = 0.13
r = –0.79
P = 1.6 × 10−9
D. longicatena log10 relative abundance (%)
Dietary fiber
SBP
Vehicle
Day
5
12
9
21
I
DSM 13814
log10(relative abundance[%])
13SM
14SM

**Figure S5. Effects of Arabinan on Hadza *D. longicatena* and Bacterial Communities in Experimental Models (Related to Figure 5)**

(A) Growth curves of the 14SM community members cultured in depleted medium supplemented with 0.5% glucose, L-arabinose, or arabinan. Shaded areas indicate the standard deviation (n = 3 per group). *Clostridium spiroforme* and *F. prausnitzii* failed to grow with glucose, likely because they require additional nutrients not provided by the basal medium.

(B) Bray-Curtis distances between the 14SM plus DSM 13814 and 14SM plus ST1L communities under three carbohydrate conditions *in vitro*.

(C) Heatmap comparing the relative abundance of each community member between two carbohydrate conditions. Red and blue indicate enrichment and depletion, respectively, in arabinan-supplemented medium relative to the comparison condition.

(D) Growth curves of Hadza *D. longicatena* strain ST1L and the type strain DSM 13814 cultured in depleted medium supplemented with 10% SBP (v/v). Shaded areas indicate the standard deviation (n = 3 per group).

(E) PCoA plot of Bray-Curtis distances among fecal bacterial communities from gnotobiotic mice harboring the 14SM plus Hadza *D. longicatena* strain ST1L or the type strain DSM 13814 and administered SBP or vehicle (water) (n = 4–5 per group).

(F) Relative abundances of the 14SM members and *D. longicatena* strains in the gnotobiotic mice. Black and gray asterisks denote comparisons between the vehicle and SBP groups within the 14SM plus ST1L and 14SM plus DSM 13814 communities, respectively.

(G) Heatmaps comparing the relative abundance of each community member between the two dietary-fiber groups (left) and between the two *D. longicatena* strain groups (middle). Significant interactions between dietary fiber and *D. longicatena* strain over the course of the experiment are shown on the right. Red and blue indicate enrichment and depletion, respectively. The scale is capped at 3 and −3 to facilitate visualization of small but significant differences.

(H) Correlation between the relative abundances of *D. longicatena* and *C. scindens* in the gnotobiotic mice. The solid line and shaded area indicate the regression line and its 95% confidence interval, respectively.

(I) $Log_{10}$-transformed relative abundance of *D. longicatena* DSM 13814 in gnotobiotic mice harboring the 13SM or 14SM community, assessed seven or five days after oral administration, respectively.

One-way ANOVA followed by Tukey's post-hoc analysis (B), linear regression (C), mixed-design repeated-measures ANOVA of $log_{10}$-transformed relative abundances followed by pairwise comparisons of estimated marginal means with Holm adjustment (F and G), and Pearson's correlation (H) were used for statistical analysis. *adjusted $P < 0.05$; **adjusted $P < 0.01$; ***adjusted $P < 0.001$.
